# Portable, multilocus DNA barcoding across the diversity of meiofauna

**DOI:** 10.64898/2026.05.20.726206

**Authors:** Dylan Keene, Srishti Arya, Ben Walker, Christopher E. Laumer

## Abstract

Molecular data have revolutionised taxonomic and ecological research on the hyperdiverse communities of aquatic benthic microinvertebrates known as meiofauna. However, reference sequence databases remain highly incomplete, with variable barcode genes or fragments studied from taxon to taxon. Furthermore, there is a typical tradeoff between universality of primers and phylogenetic resolution, with rRNA markers being robustly recoverable but failing to resolve species-level divergences, and mitochondrial markers showing the reverse trend. Here, we introduce **O**xford Nanopore **r**RNA and **C**OI **a**mplicon sequencing (OrCa-seq), a rapid, low-cost protocol for parallel long-range PCR amplification and multiplexed sequencing of four amplicons, spanning the nearly-complete rRNA cistron (∼7-8 kb) and the widely studied Folmer region of COI (represented as overlapping 313 and 658 bp amplicons). This protocol, with its associated bioinformatic workflow, was designed for conducting biodiversity inventories of meiofauna and can be easily carried out in field research and educational contexts, with data available from 96-well plates of specimens within a day of lysis. To validate the method, we processed six plates of student-isolated freshwater and limno-terrestrial meiofauna, characterising the recovery of target genes and taxa with both automated and human-curated BLAST database comparisons. These data demonstrate the universal applicability of OrCa-seq across effectively all meiofauna, including the very smallest species. Nonetheless, recovery efficiency for each amplicon shows variation by taxon, with the full-length Folmer COI amplicon standing out as the most challenging. We present exemplar phylogenetic trees integrating reference sequences, demonstrating the utility of these data in confirming morphological determinations and in identifying anonymous specimens in a reverse taxonomy context. While developed in a specific educational context for use on meiofauna, the OrCa-seq approach should be readily scalable to larger research datasets, adaptable to many specimen types, and to any combination of taxon-or target-specific primers. As such, it represents a compelling multi-locus extension to the ever-growing repertoire of nanopore DNA barcoding protocols.

## Introduction

Meiofauna are the microscopic, motile invertebrate fauna associated with aquatic sediments, vegetation, and leaf litter & soils (in which environments they are sometimes considered a part of the “mesofauna”). These communities play essential ecological roles, e.g. in nutrient cycling as an interface between microbes and larger animals, and in structuring the sedimentary environment (Schratzberger and Ingels, 2018). A large fraction of aquatic biodiversity is also represented in this size fraction, and the phylogenetic diversity of meiofauna is unparalleled, with ∼⅔ of animal phyla represented (Blaxter, 2016; Worsaae et al., 2023).

This high diversity and their typically short lifecycles also posit meiofauna communities as promising sentinels of environmental change (Martínez et al., 2025a, 2025b; Moens et al., 2022). However, this high diversity also makes comprehensive ecological studies challenging, as morphological identification to the species level is labour-intensive, limited by microscopy/histology/library resources, and often requires years of taxonomic expertise. Consequently, ecologists have historically restricted focus to a small number of abundant taxa (often, nematodes), and limited identification to higher taxa such as families or genera.

In this light, molecular methods have been transformative for meiofauna research, albeit not without their own challenges. DNA barcoding of individual specimens has been widely used for primary identification (Hebert et al., 2003), with a common outcome being the recognition of pervasive cryptic speciation (Fontaneto et al., 2009; Leasi and Norenburg, 2014; Tessens et al., 2021). The ribosomal RNAs (rRNAs) contain highly conserved regions enabling the design of true universal primers. For many taxa, 18S, and less commonly 28S, has hence been adopted as the *de facto* barcode, not least since these genes were the first to be deposited in sequence databases, leading to a positive feedback loop in marker desirability. The widespread availability of rRNAs on GenBank, and their overall gradual evolutionary rate, has motivated innovative approaches such as “reverse taxonomy”, in which large numbers of effectively anonymous meiofauna are rRNA-sequenced and post-hoc assigned to putative taxa, representing a compelling approach to biodiversity discovery in such phylogenetically disparate communities (Markmann and Tautz, 2005)

However, such slow-evolving markers are limited in their ability to discriminate related species (Tang et al., 2012), especially when studying amplicons within the Sanger (<800 bp) or short-read (<300 bp) optimal fragment size range. Fast markers such as mitochondrial COI, favoured in e.g. insect barcoding, should theoretically solve this problem, but the rapid substitution rates and high phylogenetic spread represented in the meiofauna challenge the proposed universality of widely-used degenerate primer pairs. Many orders or even families, such as within Platyhelminthes, require taxon-specific redesign of primers (Cuadrado et al., 2026; Vanhove et al., 2013) for successful barcoding. An independent problem - contamination with gut contents, symbionts, and environmental DNA - stems from small body size, necessitating the use of whole specimens for DNA extraction. While reflecting real ecological relationships (Maghsoud et al., 2014), such contaminants remain a practical problem in DNA taxonomy, as degenerate primers can preferentially amplify off-target DNA or necessitate cloning to resolve mixed templates when using Sanger sequencing.

Third-generation sequencing offers an opportunity to solve many of these problems. The length of amplicons suitable for these technologies is limited only by template integrity and what the chosen polymerase can successfully amplify. For instance, amplicons spanning full-length 18S, ITS-1, 5.8S, ITS-2, and 28S rRNA genes are well within reach (Krehenwinkel et al., 2019). Because true single molecules are read, mixed templates can be resolved as separate molecular Operational Taxonomic Units (mOTUs) with read clustering and consensus formation, or as amplicon sequence variants (ASVs) (Riisgaard-Jensen et al., 2026; Vierstraete and Braeckman, 2022). Both PacBio and Oxford Nanopore platforms have improved dramatically in throughput recently, with 10,000-100,000 barcodes now achievable within a single sequencing run (Hebert et al., 2025, 2018).

Nanopore sequencing is particularly unique in its portability and ease of sample preparation, with amplicon sequencing demonstrated in rudimentary field labs (Krehenwinkel et al., 2019), and with realistic specimen-to-sequence turnarounds of less than a working day (Vasilita et al., 2024).

As part of a training workshop on the natural history and taxonomy of freshwater and limno-terrestrial meiofauna (Majdi et al., 2026), we leveraged this potential to incorporate a DNA taxonomy module into our activities. The protocol we developed for this purpose, OrCa-seq (**O**xford Nanopore **r**RNA and **C**OI **A**mplicon **seq**uencing), takes a 96-well plate of isolated meiofauna specimens from extraction to sequencing in under 24 hours, representing nearly the full-length rRNA cistron excluding external transcribed spacers, plus the “Folmer” fragment of COI, by co-indexing four amplicons in multiplex (Fig. 1A). We validate this approach on six plates of student-isolated specimens. This dataset spans 11 phylum or class-level higher taxa, including some of the very smallest meiofauna (e.g. chaetonotoid gastrotrichs, monogonont rotifers, and single ciliophoran cells).

**Fig. 1.**
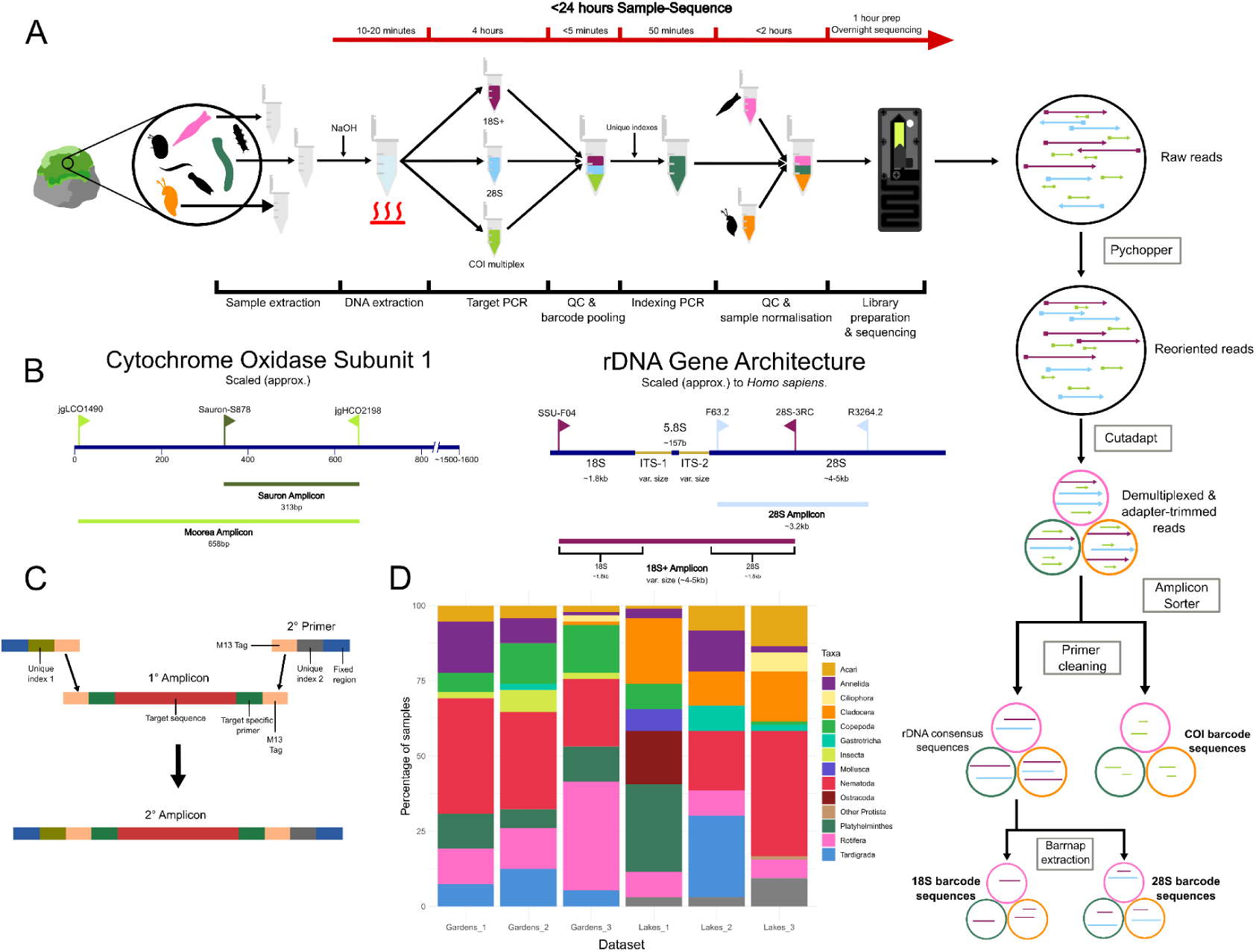
ONT-based rDNA & COI Amplicon sequencing (ORCA-seq): Workflow, molecular rationale, and collection data. A: End-to-end laboratory (L-R) and bioinformatic (T-B) workflow for ORCA-seq, time course approximate. B: Gene architecture for ORCA-seq targets, with locations of primer binding sites. Below each gene, approx. sizes and given names of the PCR fragments generated are shown. All scales are approximate. C: Molecular mechanism for the barcode-agnostic, sample-specific dual-combination indexing used in ORCA-seq. Not to scale. 1° Amplicon is any of the amplicons generated in one of the three target PCRs. 2° Primer is the Indexing primer. D: Stacked barplot of meio & macro faunal taxa sampled as part of each of the six datasets studied here. “Gardens” datasets collected in the NHM gardens, London, UK. “Lakes” datasets collected in the Lake District National Park, UK.

## Methods

### Specimen collection

Soil, leaf litter, moss, and pond sediment were collected in the Natural History Museum’s wildlife garden in June 2025. Meiofaunal extraction was undertaken with an adaptation of the Whitehead-Hemming tray method (Whitehead and Hemming, 1965) using seed-sprouting trays. For waterlogged samples, we used the oxygen-depletion method (Schockaert, 1996). Reverse taxonomy specimens were isolated from petri dishes and were identified only to phylum or class at the stereomicroscope (Brunel, BMZE). Those given a morphological identification prior to lysis were examined with a Leica DM500 compound microscope (or similar). We isolated specimens into the bottom of 8-tube PCR strips in a lens of 1-2 µL of bottled spring water, or ambient water passed through a sterilizing (0.2 µm) filter.

### Tissue lysis

Instead of full DNA extraction, we adapted the Hot Sodium Hydroxide and Tris (HotSHOT) protocol (Truett et al., 2000) to obtain crude, PCR-amplifiable lysates, a strategy that had previously been shown to be compatible with meiofauna barcoding (Amezcua-Martínez et al., 2025). We prepared an alkaline lysis buffer (25 mM NaOH, 0.2 mM EDTA, pH 12.0), preheated to 95°C immediately before use. 25 µL of hot lysis buffer was added to each tube before incubation at 95°C for 10 minutes. Animals with a sclerotised cuticle were lysed for up to 20 minutes. Post-lysis, a neutralisation buffer consisting of 40mM Tris-HCl (pH 5.0) was added in equal volumes, and mixed by vortexing. To limit DNA degradation, samples were stored on ice or at-20°C when not in use.

### Target gene amplification

Per sample, three PCR reactions were carried out to amplify four separate amplicons from within either the mitochondrial gene COI or the nuclear ribosomal repeats (Fig. 1B). All target primer sequences were adapted from their original sequences by 5’ appending them with short M13-derived tags (M13F: TGTAAAACGACGGCCAG, M13R: CAGGAAACAGCTATGAC) (Ivanova et al., 2007). Specific primer sequences are in Supplementary Table 1. Primer-pair universality and PCR conditions were initially validated using 6 model freshwater invertebrate species (*Caenorhabditis elegans* N2, *Adineta vaga* AD008, *Hypsibius exemplaris* Z151, *Lepidodermella squamata*, *Pristina leidyi*, and *Stenostomum constrictum*) kept in culture at the NHM (data not shown). PCR master mixes were constructed as per manufacturer recommendations but cut to 12.5 µL total volume, with 2 µL of crude DNA extract as input template.

The 4-5 kbp amplicon termed “18S+”, spanning ∼1800 bp of 18S, the complete ITS1-5.8S-ITS-2 segment, and ∼1800 bp of 28S rRNA, was used with forward primer SSU_F04 (Blaxter et al., 1998) and reverse primer 28S_3RC (Machida and Knowlton, 2012) at 300 nM each, with 2 µl of DNA lysate as input. The 3.2 kbp “28S” amplicon, completely overlapping the segment of 28S rRNA in 18S+, used forward primer F63.2 and reverse primer R3264.2 at 300 nM each (Passamaneck et al., 2004). Both the 18S+ and 28S reactions utilised Terra™ Direct PCR Polymerase Mix (TaKaRa Bio cat. no. 639271) with 0.125 µL polymerase per reaction, and were run according to the same protocol: 1x[98°C/2:00], 35x[98°C/0:10, 62°C/0:15, 68°C/6:00], 1x[68°C/5:00].

Two overlapping fragments of the COI gene were amplified using One*Taq*™ Hot Start DNA polymerase (New England Biolabs, cat. M0481) in a single multiplex reaction. Forward primers, jgLCO1490 (Geller et al., 2013) and Sauron-S878 (Rennstam Rubbmark et al., 2018) were at 200nM and 100nM concentration, respectively, with the reverse primer shared by both amplicons, jgHCO2198, at 300nM. These conditions co-amplify a 313 bp amplicon (“Sauron”) and a 658 bp amplicon (“Moorea”), expected to give greater and lesser taxonomic universality, respectively. The PCR conditions are as follows for all Gardens samples: 1x[94°C/2:00], 40x[94°C/0:15, 54°C/0:15, 68°C/1:00], 1x[68°C/5:00]. Whilst in-field (‘Lakes’ samples) annealing temperature was lowered to 47°C for plates 2 and 3.

### Quality control, indexing PCR, sequencing library preparation

PCR products were assessed through gel electrophoresis of all individual reaction products. These were run on 1.1% w/v Agarose in TAE gels, with 1x Diamond™ Nucleic Acid Dye (Promega, cat. H1181) as a component of the loading buffer. Samples were run alongside Hyperladder™ 1kb (Meridian Bioscience BIO-33025). At the NHM, gels were visualised on the Syngene G-Box. In the Lake District, a BioRad GelDoc Go was used for this purpose.

The PCR products were pooled without cleanup with 1.5, 1.5, and 1 µL taken respectively from of reactions 18S+, 28S, and COI. 1 µL per barcode pool was then taken forward for a second round of indexing PCR intended to co-index all 4 amplicons, using 12.5 µL One*Taq*™ Hot Start DNA polymerase (Fig. 1C). We designed bespoke indexing primers constituting the Element Biosciences SP5 or SP27 tag, a 17 bp indel-minimizing barcode designed using Freebarcodes (Hawkins et al., 2018), and the M13F/R sequence, in 5’ to 3’ orientation (supp. Table 1 for sequences). Amplicon pools were given unique single indexing primer combinations, with each primer at 200nM. PCR conditions were as follows: 1x[94°C/2:00], 7x[94°C/0:15, 47°C/0:15, 68°C/4:00], 1x[68°C/5:00]. Cycles were limited to seven to avoid overamplification.

Indexed samples were pooled in groups of 8, cleaned up using a DIY SPRI recipe (Rohland and Reich, 2012) at a 0.8X ratio of bead suspension: PCR master mix, and quantified with the Qubit™ dsDNA HS kit (ThermoFisher cat. Q32851). For each sequencing run, 96x samples (12x pools of the 8-specimen groups) were normalised to the lowest molecular weight in the batch, to generate a pool containing 100-200 fmol.

Library preparation was using the ONT ligation sequencing kit SQK-LSK114 with minor alterations. The standard protocol was abridged to begin at adapter ligation, due to the A-tailing ability of the indexing polymerase. Ligation of the nanopore adapter was carried out at typically ¼ the standard volume of the R10.4.1 “Ligation” kit (Oxford Nanopore Technologies, cat. LSK114). ‘Gardens’ plate 1 was loaded onto a MinION flow cell and run on a GridION, whilst 2 & 3 were loaded onto PromethION flow cells and run on a P2 Solo for an average of 1hr 47min (MinKNOW v25.05.14), basecalling using “SUP” accuracy. All ‘Lakes’ samples were loaded onto a single flow cell on a MinION Mk1D, basecalled to “HAC”, and allowed to sequence for an average of 12 hr 49 min (Supplementary Table 2).

Between each run, the flow cell was washed with the EXP-WSH004 kit.

### Bioinformatic processing of barcodes

Raw sequencing reads were processed with `PyChopper(`v2.7.0) using parameters `--k LSK114 --Q 10 --p —-m edlib` (Sipos et al., 2022) to reorient cDNA reads to a consistent sense orientation based on adapter positioning. Reoriented reads were then demultiplexed using `Cutadapt (`v4.9.0) with parameters `–action=trim --e 0.1 –-rc.` Forward and reverse primers were denoted using `-g and-a` flags, respectively (Martin, 2011). Only bins corresponding to valid adapter combinations were retained for downstream analysis.

`AmpliconSorter` with the `-ar` option (Vierstraete and Braeckman, 2022) was used to sort reads by length (COI amplicons: 300–900 bp; rDNA amplicons: >2,000 bp) and to cluster reads into consensus sequences using the default identity threshold. Primer sequences were trimmed from consensus amplicons using `Cutadapt`: linked primer pairs were removed simultaneously (`-g fwd…rev`). Residual primer contamination was assessed by searching sequences with `Seqkit subseq-r 1:100` (Shen et al., 2016); any sequence retaining a primer match was discarded.

COI sequences required no additional processing. For rDNA amplicons, `Barrnap v0.9` (Seemann, 2025) was used to annotate 18S and 28S genes; 5.8S sequences were discarded. For submission to NCBI, `pyBarrnap v0.5.1` (Shimoyama, 2024) with `--accurate` mode was used instead of `barrnap` to more precisely delineate boundaries between the 5.8S, ITS2, and 28S regions of sequences derived from the 18S+ amplicon.

Computational analyses were performed on Crop Diversity HPC, described by (Percival-Alwyn et al., 2025).

### Automated BLAST searching

Command-line `BLASTn v2.16.0` (Altschul et al., 1990) with blastdb v5 (25th May, 2025) was run on all gene sequences from all barcodes, requesting 500 target alignments, and selecting the top 5 of these based on e-value. The lowest common ancestor for each contig was calculated in `R v4.4.2` by retrieving taxonomy ranks using `taxonomizr v0.11.1`, and selecting the lowest common rank of the 5 highest e-value BLAST hits per contig. For both automated and manual BLAST searches, we discarded results from non-meiofaunal and low-abundance taxa in the dataset (Mollusca, Insecta and other Protista) before analysis.

### Manual BLAST hit curation

For each barcode (18S, 28S, Moorea, Sauron), fasta files of the maximum readcount contig per sample were compiled, and webBLAST blastn run against NR (November 2025–January 2026). Hits were manually evaluated by e-value, bit score, percent identity, and query cover, and the best taxonomic assignment from the top 5 results was recovered, excluding anonymous environmental isolates. This curation accounted for cases such as high-identity matches outranked by higher-bitscore hits or likely misidentified sequences. Assignments were compared against expected specimen taxa and given a pass/fail result. Where the max readcount contig failed, a command-line BLAST database from the remaining consensus sequences was built for that sample and queried with a taxon-specific reference sequence (Supplementary Table 3). Hits were confirmed via webBLAST as above. For Sauron and Moorea amplicons, tblastx was additionally used when blastn returned no strong hit, in both command-line and web searches.

Data was analysed in R, and visualised using ggplot v4.0.2 and Inkscape v1.4.2.

### Maximum likelihood treebuilding

Best-matching contigs were filtered and grouped by higher taxon using the custom script `08_reorganise_barcodes_per_taxon.sh`. For phylogenetic placement of sample barcodes, a maximum of 5000 representative reference sequences per gene were mined from NCBI using `Gene-Fetch v1.0.20` (Parsons and Price, 2025), setting minimum length as 100bp. This was completed for Nematode 18S and Annelid COI. For Platyhelminth 18S, a maximum of 500 sequences for each turbellarian order were retrieved (excluding Neodermata). `MAFFT` (v7.526) `--auto` (Katoh and Standley, 2013) was used to globally align all reference and sample sequences. A per-taxon and per-barcode distance matrix was computed with `R v4.5.3`, Ape v5.8.1 (Paradis et al., 2004), and Phangorn v2.12.1 (Schliep, 2011)with the custom script `phylo_anchor_filter.Rmd`. References were filtered by excluding those with >3*median divergence from any sample, white-listing those meeting a minimum similarity threshold, and de-duplicating to retain 3 per similarity group (Supplementary Table 4).

De-duplicated white-listed references and non-white-listed references were subset, selected from a `FastTree v2.2.0` (Price et al., 2009) maximum-likelihood tree to maximise phylogenetic diversity (PD) using Faith’s PD metrics (Table 4). Further manual tip filtering was then completed, and samples and reference sequences were re-aligned.

Maximum-likelihood trees were built with `IQ-TREE v2.3.6` (Minh et al., 2020) using `-mfp`,`-B 1000`, and `-bnni` for model finding and bootstrapping. `TreeViewer V2.2.0` was used for visualisation and wrangling (Bianchini and Sánchez-Baracaldo, 2024).

Tips in each phylogenetic tree were manually inspected for misidentification.

Sequences appearing in anomalous positions given their assigned taxonomic identification were searched against nr with webBLASTn and highlighted. The closest hit, its percentage identity, accessions, and the lowest common ancestor of the top 5 hits were noted in Table S8.

### AI usage statement

AI (claude.ai sonnet v4.5) was used to assist with writing and troubleshooting data processing, analysis and visualisation code, as well as composing CO1_splitter_maxread_extractor.py and ribo_maxread_extractor.py.

## Results

### Oxford Nanopore rRNA and COI amplicon sequencing (OrCA-seq)

We collected and barcoded individual 557 specimens in two batches (Fig. 1D).

Initially, 284 specimens were collected from the NHM’s wildlife gardens (“Gardens” samples) in a reverse taxonomy fashion, identifying specimens only to phylum or class. We then collected a further 273 specimens as part of a meiofauna systematics field course in the Lake District, United Kingdom (“Lakes” samples). Students, working in collaboration with taxon-specific experts, identified specimens to the lowest taxonomic rank possible before barcoding. These were split into three 96-well plates per location, and contained a diverse representation of meiofaunal taxa (Supplementary Fig. 1).

We processed each plate in a separate OrCA-seq run, co-indexing all four amplicons. The mean output across all runs was 2.39Gb, with 1.77M reads, and a mean N50 of 3.27kb (Supplementary Table 2); read Phred score averages were typically between 15-20 (Supplementary Fig. 2A-F). Post-sequencing, we extracted barcodes with a custom workflow, by quality-controlling reads, demultiplexing them, and clustering them into consensus sequences, before removing primers and annotating rRNA gene boundaries (Fig. 1A, Supplementary Fig. 2G). Anywhere between 0 and 30 contigs were retrieved post-clustering (Supplementary Fig. 2H).

### Assembly and automated taxonomic assignment of consensus sequences

OrCA-seq recovered thousands of distinct amplicon consensus sequences per gene from the 557 specimens we attempted to barcode, nearly all of which show tens or hundreds of contributing reads (Fig 2A, Supplementary Table 5). The 18S+ and 28S amplicons recovered broadly similar numbers of contigs (median 10 per sample), whereas from COI, the Moorea amplicon represented the specimens with both lower average coverage and circa 1/2 as many contigs as the Sauron amplicon (medians 7 and 15, respectively).

**Fig. 2.**
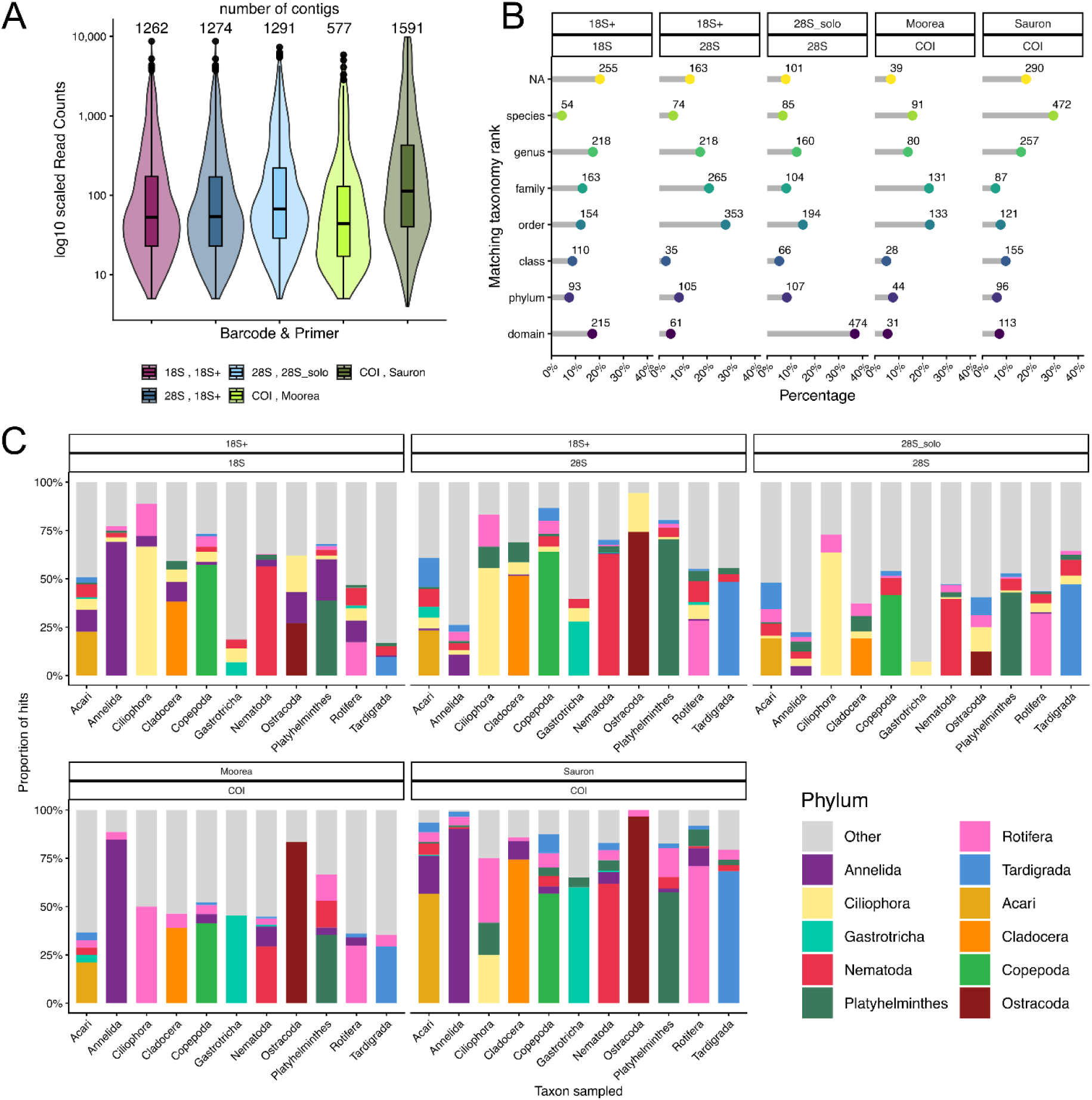
Broad assessment and taxonomy of barcode sequences. (A) Violin plot of log10-scaled read counts per contig, split by amplicon. Total contigs per barcode listed above violins. (B) Lollipop plots summarising percentages of taxonomic rank of last common ancestors for each contig per barcode and primer pair, ranked by taxonomic level. Absolute counts of contigs for each category listed by each lollipop. NA assigned if no last common ancestor could be computed. (C) Total proportions of contigs matching each taxon intentionally sampled. Other/NA group assigned if hits are to unintentionally sampled taxa or are unidentifiable.

We next assessed the accuracy of OrCA-seq barcodes for taxonomic assignment relative to taxa intentionally sampled through BLAST searching against NR and calculating the last common ancestor. rRNA barcodes less frequently achieved species-level assignment compared to COI barcodes (Fig. 2B). A substantial proportion of contigs either could not be assigned or were resolved only to the domain level, consistent with documented limitations of LCA-based taxonomic assignment (Fosso et al., 2018).

In aggregate, we observe that the plurality of hits in nr match the expected higher taxon for most genes and for most taxa (Fig 2C, Supplementary Fig. 3). Nearly all taxa also show many off-target hits, often including other meiofaunal taxa, with proportionately fewer on-target hits from the ribosomal than the COI marker.

### Curated assessment of taxon-specific barcode retrieval

As meiofaunal taxa often show limited GenBank representation (Martínez et al., 2025b) and anonymous, misidentified, and variable-length sequences are frequent, we assessed target taxa representation by OrCA-seq barcodes specimen-by-specimen, using webBLAST against the NR database. We expected a maximum of 2152 contigs total, assuming one per amplicon per specimen (18S, 28S, Moorea, Sauron). Of these, 27.04% (582) had no assembled sequences representing that gene, and 19.70% (424) matched only non-targeted taxa during the manual BLAST curation. Plausible hits were found in 53.25% (1146) of assessed markers, with the majority (1031, 47.91% of total contigs) originating from the contig within each sample barcode file with the highest readcount associated, the “Maximum Readcount Contig” (MRC). A smaller percentage of matches (5.34% of total contigs) arose from an “Alternate Contig” (AC), defined as a sequence showing a convincing match to the targeted taxon, albeit with less than the maximal readcount from that specimen and amplicon.

As expected from the high conservation of priming sites in nuclear rRNAs, these genes give the highest success rates (Fig. 3A). For 18S rRNA, we retrieved targets from 63.75% (91.55% MRC, 8.45% AC) of potential contigs, with 21.93% of contigs missing and 14.31% off target. The recovery success rate was the highest for 28S rRNA, with 77.32% (91.83% MRC, 8.17% AC) of potential contigs yielding positive matches, with 12.64% missing and 10.04% off target. COI had the lowest overall percentage of positive BLAST identification, with only 35.97% (86.56% MRC, 13.43% AC) of potential contigs leading to a positive identification. 36.80% of potential contigs were missing, and 27.23% were off-target matches.

**Fig. 3.**
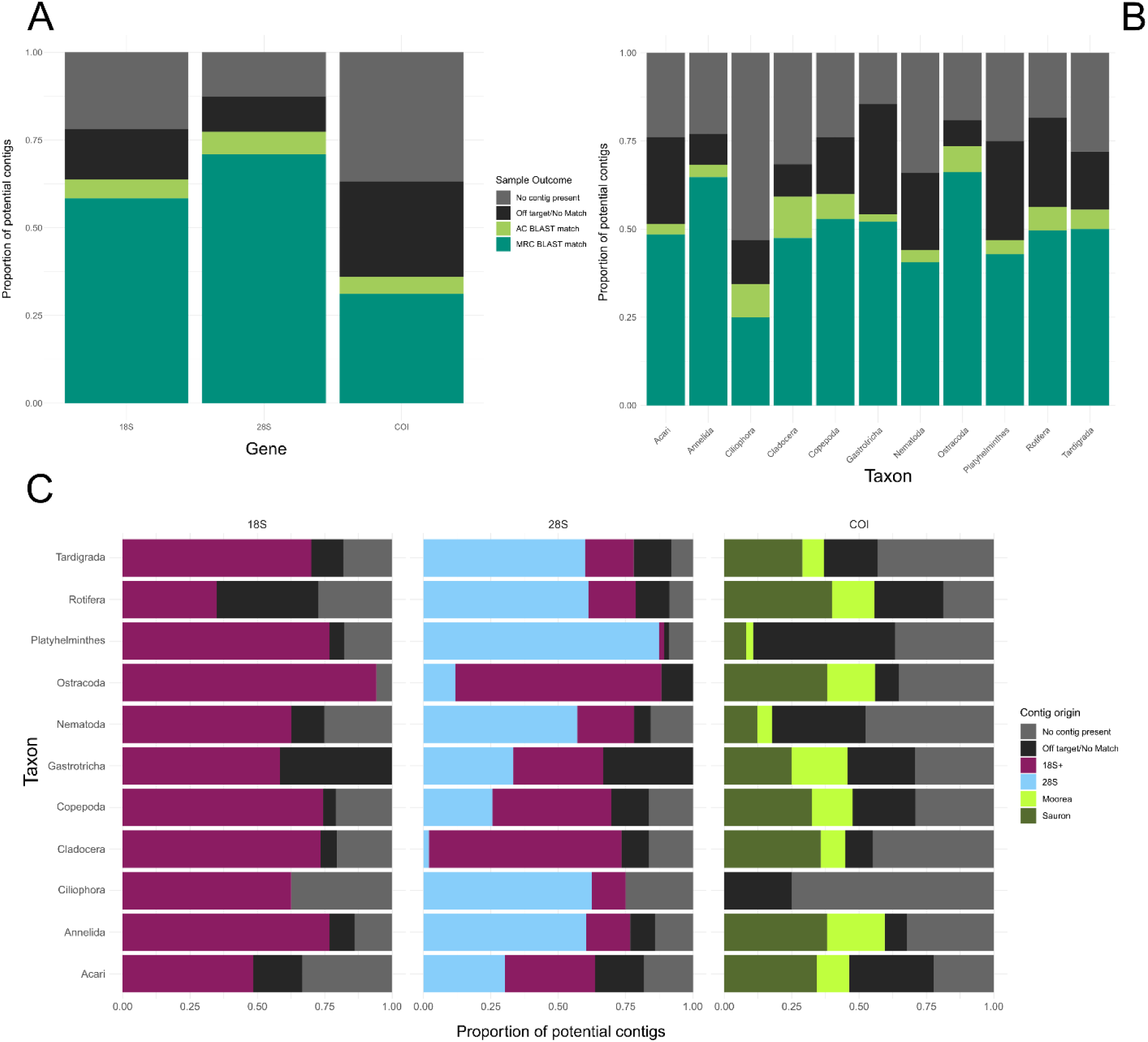
Assessment of barcoding success from the manual BLAST curation of contigs. A: Stacked barplot of the proportion of all meiofaunal samples attaining a positive ID from assembled contigs. Split by gene. Proportions required as the COI column contains both Moorea and Sauron BLAST results. 28S contains both 18S+ and 28S derived amplicons. The max readcount contig is the sequence for each sample with the highest readcount associated with it by AmpliconSorter. Alternate contigs are only assessed if the max readcount contig does not match the expected taxonomic group. B: Stacked barplot of the proportion of all meiofaunal samples attaining a positive ID from assembled contigs. Split by taxa. Key shared with panel A. C: Stacked barplot of the outcome of manual BLAST curation by meiofaunal taxa, split by barcode, coloured by amplicon of origin for each.

Aggregating by target taxon, we see substantial clade-by-clade variability in the success rate for each (Fig. 3B). Across all barcodes (Fig. 3B, Supp. Table 6), Ostracoda showed the highest recovery rate (73.53%), followed by Annelida (68.24%), Copepoda (59.88%), Cladocera (59.18%), Rotifera (56.25%), Tardigrada (55.50%), Gastrotricha (54.17%), Acari (51.49%), Platyhelminthes (46.88%), Nematoda (44.05%) and Ciliophora (34.38%) (Supplementary Table 6). Taxon-specific recovery rates for 18S rRNA ranged from 35% (Rotifera) to 94.12% (Ostracoda).

The recovery rate of 28S is high overall, and less variable than 18S (the maximum difference in recovery being 25.65%), partly because it combines results from the 3.2 kb 28S amplicon as well as the 1.8 kb fragment of this gene from the “18S+” amplicon. However, there are substantial taxonomic differences in which amplicon represents 28S. We observed amplicon bias in these results through relative percentages of 18S+ and 28S amplicons seen in successfully recovered contigs (Fig 3.C). We saw the most 28S bias within Platyhelminthes (98% 28S), with Ciliophora (83.33%), Annelida (78.79%), Rotifera (77.78%), Tardigrada (76.92%) and Nematoda (73.04%) all being 28S-biased (Fig 3.C, Sup. Table 7.B). Gastrotricha was evenly split between the two amplicons. The 18S+ preferred taxa all belonged to phylum Arthropoda. An uneven distribution is also seen when comparing the amplicon of origin ratio within the successfully identified COI gene contigs (Moorea: Sauron). Across all taxa (discounting Ciliophora, in which no contigs were successfully identified), the majority of positively identified contigs were Sauron-derived, with the most even proportion in Gastrotricha samples (54.55% Sauron) and the most Sauron-biased in Cladocera (79.55% Sauron) (Supplementary Table 7.C).

### Reliable phylogenetic placement of barcoded specimens

To demonstrate the utility of OrCA-seq, we built maximum-likelihood phylogenies for three of the most widely sequenced phyla (Fig. 4), mining reference sequences within a pairwise sequence similarity threshold of our novel barcodes. These trees combine specimens identified to the lowest possible level, typically species (Lakes data), as well as anonymous specimens (Gardens data).

**Fig. 4:**
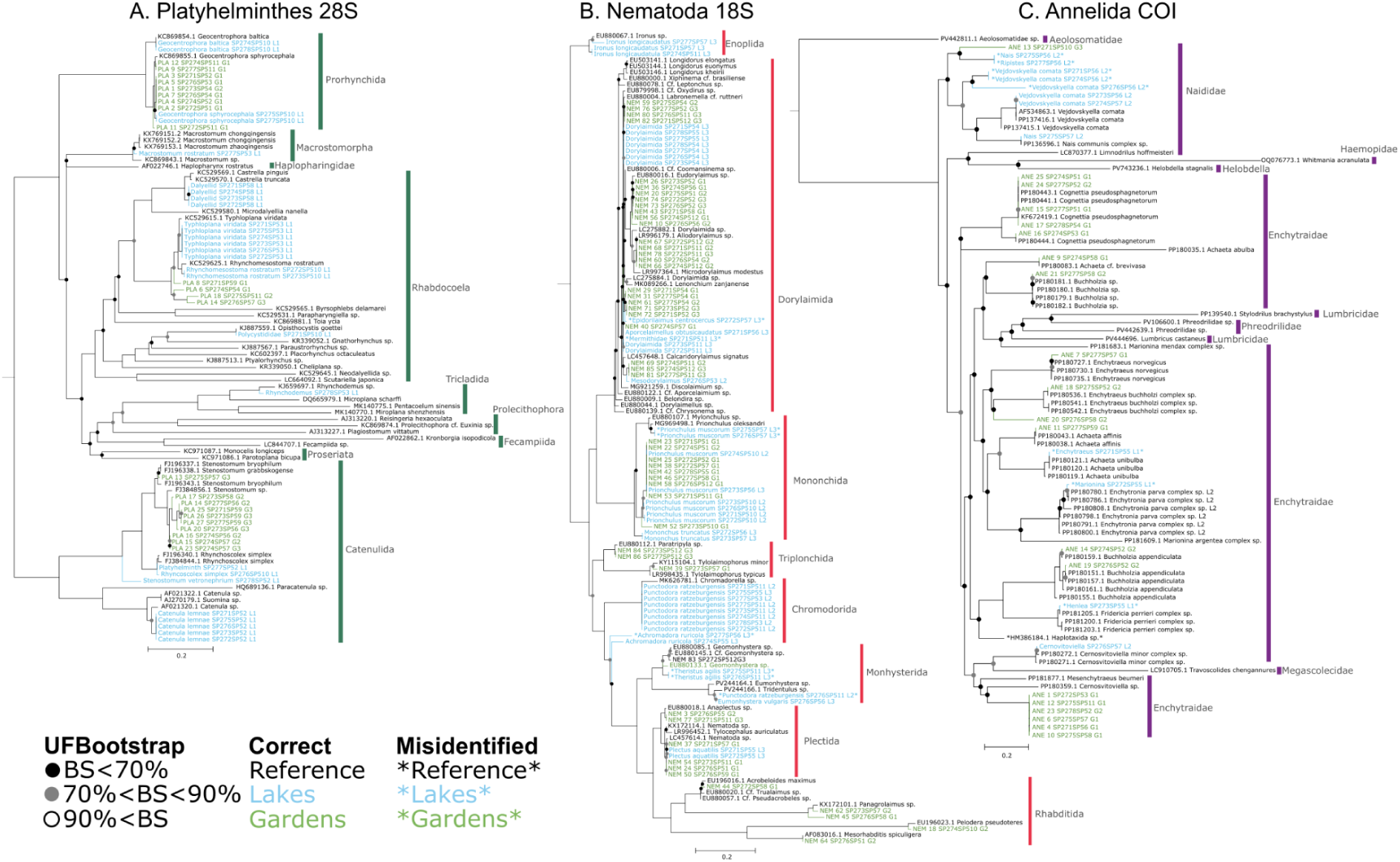
Maximum-likelihood phylogenetic trees, with bootstrap scores as coloured circles on nodes, tips coloured in green for samples belonging to the Gardens dataset, tips coloured in blue for samples belonging to the Lakes dataset, and tips coloured black for reference sequences. Putatively misidentified sample and reference tips are denoted using asterisks. (A) 28S barcodes belonging to Platyhelminthes, with ML model GTR+F+R5 (B) 18S barcodes belonging to Nematodes, with ML model GTR+F+R4 (C) COI barcodes belonging to Annelids, with Moorea primer and Sauron primer amplified barcodes for the same samples collapsed, and ML model GTR+F+I+G4.

In the platyhelminth 28S tree, sampling was enriched in Catenulida, Prorhynchida and Rhabdocoela (Fig. 4A), reflecting the abundance of these microturbellarian orders in freshwaters (Fig. 4A). The topology confirms the identity of many specimens as widely distributed species known to occur in the UK, such as *Rhynchomesostoma rostratum*, *Opisthocystis goetti*, and *Rhynchoscolex simplex*. Six specimens assigned to *Typhloplana viridata* collectively show 2% divergence (in pairwise blastn) from the reference sequence but not within samples, plausibly representing cryptic speciation. This is also the case for *Geocentrophora sphyrocephala.* Additionally, we see evidence for other species either not yet represented in GenBank by 28S, and/or new to science, including another 4 anonymous sequences of Typhloplanidae (Rhabdocoela), and several Clade IV Stenostomidae (Tratkiewicz et al., 2026).

In the nematode 18S tree (Fig. 4B), we see broad phylogenetic sampling across 8 orders, reflecting this phylum’s frequent ecological transitions to freshwater (Holterman et al., 2019). Some evidence of cryptic species widely distributed in the UK can be seen, e.g., in specimens assigned to *Prionchulus muscorum*. Near-or total-identity matches between many of the anonymous nematodes and reference 18S sequences, e.g., *Mesorhabditis spiculigera* and *Allodorylaimus* sp., confirm the utility of OrCA-seq in a reverse-taxonomy context.

In the annelid COI tree, individuals broadly distributed among Clitellata are represented (Fig. 4C), with Enchytraeidae and Naididae representing the best-sampled families. Deep divergences are poorly resolved as expected for COI, e.g. with non-monophyly of Enchytraeidae and Lumbricidae. Numerous examples of near-identity of anonymous OrCA-seq specimens with reference barcodes are also visible in this tree, e.g. in *Cognettia pseudosphagnetorum*, *Achaeta affinis*, and *Buchholzia sp.* In the conventionally identified specimens, there are cases of both the morphological labels matching reference sequences, e.g. for *Cernosvitoviella* sp. and *Vejdovskyella comata*, but also of clear misidentification or cryptic speciation (e.g. a specimen labelled *Marionina* sp. falling within a clade of *Enchytriona parva*, or three specimens labelled *Vejdovskyella comata* branching more deeply within Naididae).

Indeed, throughout all trees, there are some sequences whose placement suggests misidentification (labelled in Fig. 4), as expected in the context of student-led identifications (Supplementary Table 8). Some reference specimens are potentially also misidentified (e.g. HM386184, labelled as Haplotaxida but falling within Enchytraeidae). However, the many instances in which a species sampled with multiple specimens shows no branch length variation (i.e. the same haplotype) bolster confidence in OrCA-seq consensus sequence accuracy, as well as confidence in student-led taxon determinations.

## Discussion

Molecular methods centred on amplicon sequencing, most notably metabarcoding, have been transformative to the study of meiofaunal ecology. However, a major limitation of these methods is the poor sequence representation of meiofaunal taxa in reference databases (De Santiago et al., 2025; Macher et al., 2024; Martínez et al., 2025b). A related problem is the variable choice of barcode genes made by each taxonomic community, with some taxa well represented with COI but not 18S rRNA, for instance (e.g. arthropods, annelids), and with varying regions of 28S rRNA represented for each taxon.

OrCA-seq adds to the chorus of third-generation amplicon sequencing methods (Hebert et al., 2025, 2018; Krehenwinkel et al., 2019; Vasilita et al., 2024). It is unique in being intrinsically multi-locus, co-indexing three common barcoding genes for each specimen, separating reads from each gene by length and sequence similarity (Vierstraete and Braeckman, 2022). OrCA-seq thereby balances universality and low-level taxonomic resolution. Our results show robust recovery of targeted genes, particularly the long nuclear rRNA amplicons, across effectively all Metazoa, even from single individuals of the very smallest meiofauna. While we have not specifically analysed these segments, the internal transcribed spacer regions represented in ∼64% of successful 18S+ amplicons are also expected to show useful species-level variation.

Most variation in amplicon recovery success is likely due to taxon-specific primer-site divergence (e.g. in Rotifera for the 18S+ primers, or Ciliophora for the COI primers). As a general rule, the COI gene represented the most challenging gene to recover across all taxa (Fig. 3), albeit our multiplex PCR strategy allowed many more taxa to be represented by a Sauron fragment than would have amplified with only full-length Folmer/Moorea primers (Rennstam Rubbmark et al., 2018).

OrCA-seq was developed with an educational aim in mind, and its simplicity, flexibility (any target-or taxon-specific primers can be easily adapted), cost-effectiveness, and speed render it highly useful in this context. However, because of this, the validation data we present should be seen as a minimum estimate of the throughput and barcoding efficiency possible. In particular, all of our nanopore runs were allowed to run longer than needed for logistical reasons, e.g. sequencing overnight, giving an average of 983 reads per curated on-target contig. Future iterations leveraging more sophisticated and efficient multiplexing and demultiplexing (Beeloo et al., 2025) should be able to recover OrCA-seq data from thousands of specimens with a single MinION. We also expect the student-led context of these proof-of-principle data may have had a minor impact on our results in other ways, owing to the technical difficulty of handling microscopic invertebrates, i.e. with some tubes mistakenly excluding the intended specimen, or including multiple unintended individuals.

An unexpected but unavoidable pattern within these data is the presence of many distinct sequences being recovered for most amplicons, for each specimen (5,995 in total; Fig 2A). It is comforting that in the large majority of cases, a plurality of database hits match the expected taxon in automated blast screens (Fig 2C), and that overwhelmingly during manual curation, the sequence constructed from the largest number of reads matches the expected taxon (Fig 3). It is also biologically plausible that many, perhaps most unstarved, wild-caught meiofauna specimens should represent taxonomic mixtures of DNAs, including gut contents, symbionts, and plausibly, ambient environmental DNA adsorbed onto surfaces or dissolved in the small amount of freshwater accompanying the live specimen into lysis. In this sense, perhaps all DNA barcoding done using deep-coverage, third-generation sequencing with such tiny inputs and universal primers should be considered a form of metabarcoding. However, we cannot rule out that some fraction of sequences also derive from specimen-handling or laboratory cross-contamination, PCR recombination, index-hopping, or some other artificial process. Caution should therefore be applied when applying biological significance to OrCA sequences recovered with low relative coverage.

## Data availability statement

All code, configuration files, and adapter sequence files are available from the Zenodo.org archive linked below. Raw data from the manual BLAST curation, collection information, and detailed plate maps are available in the Zenodo repository here. https://zenodo.org/records/19859797?preview=1&token=eyJhbGciOiJIUzUxMiIsImlhdCI6MTc3NzQ3MzE0MiwiZXhwIjoxNzk4Njc1MTk5fQ.eyJpZCI6IjU1MGQyMTQ3LWVlNmEtNGIzZS05ZDg0LWE2N2VlOTYzYThjNiIsImRhdGEiOnt9LCJyYW5kb20iOiI3YWRiZTVlODEyZTljNjk4M2ZiMzJhNGE2YjAxNWRmNiJ9.ZA4EFIhZiqY7ydZpnHw0eTYUCPS_Kz9Y_lm9H6GH_XUKlhUDKf-Rx6dSXpbpX2-LuNf-TZreowv-uSioIielGg.

A detailed wet-lab protocol is available at: https://www.protocols.io/blind/4A67195D43D711F1BE9F0A58A9FEAC02

## Acknowledgements

We are grateful to the students (Andrea Lenti, Clay Bibby, Erik Tihelka, Giovane Vedovatti, Hannah Budroe, Hannah King, Isobel Evans, Jarosław Brodecki, Katarzyna Tratkiewicz, Louise Thurston, Mateo Carvajal, Matylda Gajda, Vojtěch Brož, Wiktoria Dmuchowska, and Will Dawson), co-instructors (Alex Kieneke, André R.S. Garraffoni, Christer Erséus, David Horne, David Wilkinson, Diego Fontaneto, Genoveva Esteban, Harry Smit, Kay Van Damme, Matthew Shepherd, Nabil Majdi, Rony Huys, Walter Traunspurger, and Witold Morek), and other participants (Ewan Shilland, Gill Notman, and Tsvetoslav Georgiev) of the 2025 Lake District Meiofauna Workshop for their labours in collecting and identifying specimens. We also acknowledge the staff at the University of Cumbria Ambleside, in particular Joanne Jones and Helen Wright, with whose support we felt at home in the teaching labs. The NHM Molecular Biology Laboratories supported us during the initial development of OrCA-seq. All specimens were collected under permissions granted by the National Trust, the Cumbria Wildlife Trust, Forestry England, and Natural England. Ben Walker’s summer research placement was funded by the NERC IGNITE DTP, grant no. UKRI1327. Chris Laumer is funded by a Royal Society University Research Fellowship, URF\R1\221744. The workshop itself was funded by the NERC “Delivering training courses for environmental scientists 2024”, UKRI1312, with additional funding from the Royal Society’s Climate and Biodiversity Community Engagement Programme.

## Author contributions

CEL conceived the project, set its goals and scope, and designed the experimental methodology. DK and BW performed wet-lab protocol optimization and data collection. SA led on bioinformatic analysis, with contributions from CEL and DK. CEL led writing of the manuscript, with figures, supplemental material, and additional writing contributions led by DK and SA. All authors contributed critically to the drafts and gave final approval for publication.

## Statement on inclusion

This study was undertaken entirely using specimens found within the UK, where all authors reside. It was designed within the context of a training workshop that included delegates from 12 countries, who are thanked by name in the Acknowledgements.

## References

Altschul, S.F., Gish, W., Miller, W., Myers, E.W., Lipman, D.J., 1990. Basic local alignment search tool. Journal of Molecular Biology 215, 403–410. 10.1016/S0022-2836(05)80360-2

Amezcua-Martínez, C.F., Jiménez-Marín, A.R., Dueñas-Cedillo, A.R., Flores-Martínez, J.J., Gomez-Lunar, Z., Ruiz, E.A., Armendáriz-Toledano, F., 2025. HotSHOT Extraction Is a Fast, Economical, and Efficient Method for Obtaining Tardigrade gDNA. Psyche: A Journal of Entomology 2025, 8887456. 10.1155/psyc/8887456

Beeloo, R., Koerkamp, R.G., Jia, X., Broekhuizen-Stins, M.J., IJken, L. van, Broens, E.M., Zomer, A., Dutilh, B.E., 2025. Barbell Resolves Demultiplexing and Trimming Issues in Nanopore Data. 10.1101/2025.10.22.683865

Bianchini, G., Sánchez-Baracaldo, P., 2024. TreeViewer: Flexible, modular software to visualise and manipulate phylogenetic trees. Ecology and Evolution 14, e10873. 10.1002/ece3.10873

Blaxter, M., 2016. Imagining Sisyphus happy: DNA barcoding and the unnamed majority. Philos Trans R Soc Lond B Biol Sci 371, 20150329. 10.1098/rstb.2015.0329

Blaxter, M.L., De Ley, P., Garey, J.R., Liu, L.X., Scheldeman, P., Vierstraete, A., Vanfleteren, J.R., Mackey, L.Y., Dorris, M., Frisse, L.M., Vida, J.T., Thomas, W.K., 1998. A molecular evolutionary framework for the phylum Nematoda. Nature 392, 71–75. 10.1038/32160

Cuadrado, D., Machordom, A., Noreña, C., 2026. A deep-time perspective on Polycladida (Platyhelminthes) through integrated phylogenetic and molecular clock analyses. Org Divers Evol. 10.1007/s13127-026-00699-0

De Santiago, A., Pereira, T.J., Ferrero, T.J., Barnes, N., Lallias, D., Creer, S., Bik, H.M., 2025. Persistent Gaps and Errors in Reference Databases Impede Ecologically Meaningful Taxonomy Assignments in 18S rRNA Studies: A Case Study of Terrestrial and Marine Nematodes. Environmental DNA 7, e70080. 10.1002/edn3.70080

Fontaneto, D., Kaya, M., Herniou, E.A., Barraclough, T.G., 2009. Extreme levels of hidden diversity in microscopic animals (Rotifera) revealed by DNA taxonomy. Molecular Phylogenetics and Evolution 53, 182–189. 10.1016/j.ympev.2009.04.011

Fosso, B., Pesole, G., Rosselló, F., Valiente, G., 2018. Unbiased Taxonomic Annotation of Metagenomic Samples. Journal of Computational Biology 25, 348–360. 10.1089/cmb.2017.0144

Geller, J., Meyer, C., Parker, M., Hawk, H., 2013. Redesign of PCR primers for mitochondrial cytochrome c oxidase subunit I for marine invertebrates and application in all-taxa biotic surveys. Molecular Ecology Resources 13, 851–861. 10.1111/1755-0998.12138

Hawkins, J.A., Jones, S.K., Finkelstein, I.J., Press, W.H., 2018. Indel-correcting DNA barcodes for high-throughput sequencing. Proceedings of the National Academy of Sciences 115, E6217–E6226. 10.1073/pnas.1802640115

Hebert, P.D.N., Braukmann, T.W.A., Prosser, S.W.J., Ratnasingham, S., deWaard, J.R., Ivanova, N.V., Janzen, D.H., Hallwachs, W., Naik, S., Sones, J.E., Zakharov, E.V., 2018. A Sequel to Sanger: amplicon sequencing that scales. BMC Genomics 19, 219. 10.1186/s12864-018-4611-3

Hebert, P.D.N., Cywinska, A., Ball, S.L., deWaard, J.R., 2003. Biological identifications through DNA barcodes. Proc Biol Sci 270, 313–321. 10.1098/rspb.2002.2218

Hebert, P.D.N., Floyd, R., Jafarpour, S., Prosser, S.W.J., 2025. Barcode 100K Specimens: In a Single Nanopore Run. Molecular Ecology Resources 25, e14028. 10.1111/1755-0998.14028

Holterman, M., Schratzberger, M., Helder, J., 2019. Nematodes as evolutionary commuters between marine, freshwater and terrestrial habitats. Biol J Linn Soc 128, 756–767. 10.1093/biolinnean/blz107

Ivanova, N.V., Zemlak, T.S., Hanner, R.H., Hebert, P.D.N., 2007. Universal primer cocktails for fish DNA barcoding. Molecular Ecology Notes 7, 544–548. 10.1111/j.1471-8286.2007.01748.x

Katoh, K., Standley, D.M., 2013. MAFFT Multiple Sequence Alignment Software Version 7: Improvements in Performance and Usability. Mol Biol Evol 30, 772–780. 10.1093/molbev/mst010

Krehenwinkel, H., Pomerantz, A., Henderson, J.B., Kennedy, S.R., Lim, J.Y., Swamy, V., Shoobridge, J.D., Graham, N., Patel, N.H., Gillespie, R.G., Prost, S., 2019. Nanopore sequencing of long ribosomal DNA amplicons enables portable and simple biodiversity assessments with high phylogenetic resolution across broad taxonomic scale. Gigascience 8, giz006. 10.1093/gigascience/giz006

Leasi, F., Norenburg, J.L., 2014. The Necessity of DNA Taxonomy to Reveal Cryptic Diversity and Spatial Distribution of Meiofauna, with a Focus on Nemertea. PLOS ONE 9, e104385. 10.1371/journal.pone.0104385

Macher, J.-N., Martínez, A., Çakir, S., Cholley, P.-E., Christoforou, E., Curini Galletti, M., van Galen, L., García-Cobo, M., Jondelius, U., de Jong, D., Leasi, F., Lemke, M., Rubio Lopez, I., Sánchez, N., Sørensen, M.V., Todaro, M.A., Renema, W., Fontaneto, D., 2024. Enhancing metabarcoding efficiency and ecological insights through integrated taxonomy and DNA reference barcoding: A case study on beach meiofauna. Molecular Ecology Resources 24, e13997. 10.1111/1755-0998.13997

Machida, R.J., Knowlton, N., 2012. PCR Primers for Metazoan Nuclear 18S and 28S Ribosomal DNA Sequences. PLOS ONE 7, e46180. 10.1371/journal.pone.0046180

Maghsoud, H., Weiss, A., Smith III, J.P.S., Litvaitis, M.K., Fegley, S.R., 2014. Diagnostic PCR can be used to illuminate meiofaunal diets and trophic relationships. Invertebrate Biology 133, 121–127. 10.1111/ivb.12048

Majdi, N., Dawson, W., Evans, I., Thurston, L., Gajda, M., Lenti, A., Smit, H., Fontaneto, D., Kamburska, L., Dmuchowska, W., Traunspurger, W., Carvajal, M., Erseus, C., Shepherd, M., Kieneke, A., Garraffoni, A.R.S., Morek, W., Shilland, E., Tihelka, E., Broz, V., Jaroslaw Brodecki, Giovane Murilo Ribeiro de Assis Vedovatti, Bibby, C.G., Tratkiewicz, K., King, H., Keene, D., Wilkinson, D.M., Huys, R., Horne, D., Budroe, H., Notman, G., Esteban, G.F., Van Damme, K., Arya, S., Laumer, C., 2026. Lake District meiofauna Workshop records. 10.15468/ZXA8J3

Markmann, M., Tautz, D., 2005. Reverse taxonomy: an approach towards determining the diversity of meiobenthic organisms based on ribosomal RNA signature sequences. Philosophical Transactions of the Royal Society B: Biological Sciences 360, 1917–1924. 10.1098/rstb.2005.1723

Martin, M., 2011. Cutadapt removes adapter sequences from high-throughput sequencing reads. EMBnet.journal 17, 10–12. 10.14806/ej.17.1.200

Martínez, A., Bonaglia, S., Di Domenico, M., Fonseca, G., Ingels, J., Jörger, K.M., Laumer, C., Leasi, F., Zeppilli, D., Baldrighi, E., Bik, H., Cepeda, D., Curini-Galletti, M., Cutter, A.D., dos Santos, G., Fattorini, S., Frisch, D., Gollner, S., Jondelius, U., Kerbl, A., Kocot, K.M., Majdi, N., Mammola, S., Martín-Durán, J.M., Menegotto, A., Montagna, P.A., Nascimento, F.J.A., Puillandre, N., Rognant, A., Sánchez, N., Santos, I.R., Schmidt-Rhaesa, A., Schratzberger, M., Semprucci, F., Shimabukuro, M., Sommerfield, P.J., Struck, T.H., Sørensen, M.V., Wallberg, A., Worsaae, K., Yamasaki, H., Fontaneto, D., 2025a. Fundamental questions in meiofauna research highlight how small but ubiquitous animals can improve our understanding of Nature. Commun Biol 8, 449. 10.1038/s42003-025-07888-1

Martínez, A., Kohler, S., García-Cobo, M., Kurtz, M.N., Fontaneto, D., Macher, J.-N., 2025b. Meiofauna as sentinels of beach ecosystems: A quantitative review of gaps and opportunities in beach meiofauna research. Estuarine, Coastal and Shelf Science 313, 109092. 10.1016/j.ecss.2024.109092

Minh, B.Q., Schmidt, H.A., Chernomor, O., Schrempf, D., Woodhams, M.D., von Haeseler, A., Lanfear, R., 2020. IQ-TREE 2: New Models and Efficient Methods for Phylogenetic Inference in the Genomic Era. Molecular Biology and Evolution 37, 1530–1534. 10.1093/molbev/msaa015

Moens, T., Sroczynska, K., Adão, H., 2022. Meiofauna in a changing world. Ecological Indicators 138, 108769. 10.1016/j.ecolind.2022.108769

Paradis, E., Claude, J., Strimmer, K., 2004. APE: Analyses of Phylogenetics and Evolution in R language. Bioinformatics 20, 289–290. 10.1093/bioinformatics/btg412

Parsons, D.A. j, Price, B., 2025. Gene Fetch: A Python tool for sequence retrieval from GenBank across the tree of life. Journal of Open Source Software 10, 8456. 10.21105/joss.08456

Passamaneck, Y.J., Schander, C., Halanych, K.M., 2004. Investigation of molluscan phylogeny using large-subunit and small-subunit nuclear rRNA sequences. Molecular Phylogenetics and Evolution 32, 25–38. 10.1016/j.ympev.2003.12.016

Percival-Alwyn, L., Barnes, I., Clark, M.D., Cockram, J., Coffey, M.P., Jones, S., Kersey, P.J., Kidner, C.A., Kosiol, C., Li, B., Marsh, W.A., Zhou, J., Caccamo, M., Milne, I., 2025. UKCropDiversity-HPC: A collaborative high-performance computing resource approach for sustainable agriculture and biodiversity conservation. PLANTS, PEOPLE, PLANET 7, 969–977. 10.1002/ppp3.10607

Rennstam Rubbmark, O., Sint, D., Horngacher, N., Traugott, M., 2018. A broadly applicable COI primer pair and an efficient single-tube amplicon library preparation protocol for metabarcoding. Ecology and Evolution 8, 12335–12350. 10.1002/ece3.4520

Riisgaard-Jensen, M., Villanelo, S.A.R., Andersen, K.S., Kirkegaard, R., Hansen, S.H., Jiang, C., Stefansen, A.V., Thomsen, J.H.D., Nielsen, P.H., Dueholm, M.K.D., 2026. Nanopore sequencing reaches amplicon sequence variant (ASV) resolution. 10.64898/2026.02.26.708165

Rohland, N., Reich, D., 2012. Cost-effective, high-throughput DNA sequencing libraries for multiplexed target capture. Genome Res. 22, 939–946. 10.1101/gr.128124.111

Schliep, K.P., 2011. phangorn: phylogenetic analysis in R. Bioinformatics 27, 592–593. 10.1093/bioinformatics/btq706

Schockaert, E.R., 1996. Turbellarians, in: Hall, G.S. (Ed.), Methods for the Examination of Organismal Diversity in Soils and Sediments. CAB International, Wallingford, pp. 211–225.

Schratzberger, M., Ingels, J., 2018. Meiofauna matters: The roles of meiofauna in benthic ecosystems. Journal of Experimental Marine Biology and Ecology, IςIMCo, the 16th International Meiofauna Conference 502, 12–25. 10.1016/j.jembe.2017.01.007

Seemann, T., 2025. tseemann/barrnap.

Shen, W., Le, S., Li, Y., Hu, F., 2016. SeqKit: A Cross-Platform and Ultrafast Toolkit for FASTA/Q File Manipulation. PLOS ONE 11, e0163962. 10.1371/journal.pone.0163962

Shimoyama, Y., 2024. pybarrnap: Python implementation of barrnap.

Sipos, B., Horner, N., Rescheneder, P., Nicholls, S., Wright, C., Rudd, S., Parker, M., 2022. epi2me-labs/pychopper: cDNA read preprocessing [WWW Document]. URL https://github.com/epi2me-labs/pychopper (accessed 10.10.24).

Tang, C.Q., Leasi, F., Obertegger, U., Kieneke, A., Barraclough, T.G., Fontaneto, D., 2012. The widely used small subunit 18S rDNA molecule greatly underestimates true diversity in biodiversity surveys of the meiofauna. Proceedings of the National Academy of Sciences 109, 16208–16212. 10.1073/pnas.1209160109

Tessens, B., Monnens, M., Backeljau, T., Jordaens, K., Van Steenkiste, N., Breman, F.C., Smeets, K., Artois, T., 2021. Is ‘everything everywhere’? Unprecedented cryptic diversity in the cosmopolitan flatworm Gyratrix hermaphroditus. Zoologica Scripta 50, 837–851. 10.1111/zsc.12507

Tratkiewicz, K., Baczyński, J., Gąsiorowski, L., 2026. Molecular phylogeny of Catenulida (Platyhelminthes) with special focus on their diversity in Poland. Zool J Linn Soc 206, zlag003. 10.1093/zoolinnean/zlag003

Truett, G.E., Heeger, P., Mynatt, R.L., Truett, A.A., Walker, J.A., Warman, M.L., 2000. Preparation of PCR-Quality Mouse Genomic DNA with Hot Sodium Hydroxide and Tris (HotSHOT). BioTechniques 29, 52–54. 10.2144/00291bm09

Vanhove, M.P.M., Tessens, B., Schoelinck, C., Jondelius, U., Littlewood, D.T.J., Artois, T., Huyse, T., 2013. Problematic barcoding in flatworms: A case-study on monogeneans and rhabdocoels (Platyhelminthes). Zookeys 355–379. 10.3897/zookeys.365.5776

Vasilita, C., Feng, V., Hansen, A.K., Hartop, E., Srivathsan, A., Struijk, R., Meier, R., 2024. Express barcoding with NextGenPCR and MinION for species-level sorting of ecological samples. Molecular Ecology Resources 24, e13922. 10.1111/1755-0998.13922

Vierstraete, A.R., Braeckman, B.P., 2022. Amplicon_sorter: A tool for reference-free amplicon sorting based on sequence similarity and for building consensus sequences. Ecology and Evolution 12, e8603. 10.1002/ece3.8603

Whitehead, A.G., Hemming, J.R., 1965. A comparison of some quantitative methods of extracting small vermiform nematodes from soil. Annals of Applied Biology 55, 25–38. 10.1111/j.1744-7348.1965.tb07864.x

Worsaae, K., Vinther, J., Sørensen, M.V., 2023. Evolution of Bilateria from a Meiofauna Perspective—Miniaturization in the Focus, in: Giere, O., Schratzberger, M. (Eds.), New Horizons in Meiobenthos Research: Profiles, Patterns and Potentials. Springer International Publishing, Cham, pp. 1–31. 10.1007/978-3-031-21622-0_1

