## Supplementary Material for "Portable, multilocus DNA barcoding across the diversity of meiofauna"

### Supplementary Materials

### Supplementary Figures

#### Supplementary Figure 1

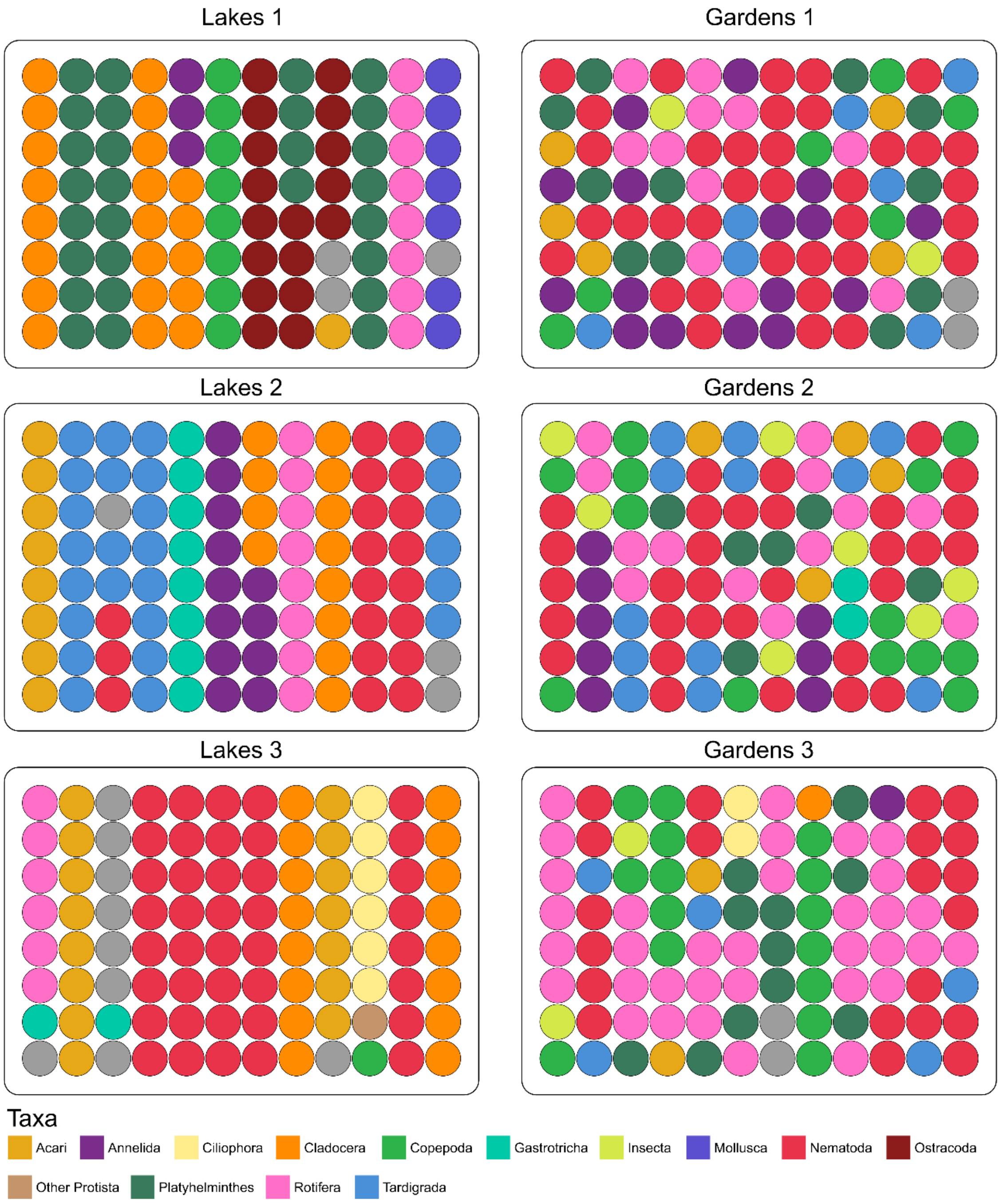

**Supplementary Fig. 1:** Taxon composition of the plates processed using OrCA-seq. “Lakes” plates were processed in the Lake District, UK using specimens collected from the surrounding areas. “Gardens” plates were collected and processed at the Natural History Museum, using specimens collected from the attached wildlife garden.

#### Supplementary Figure 2

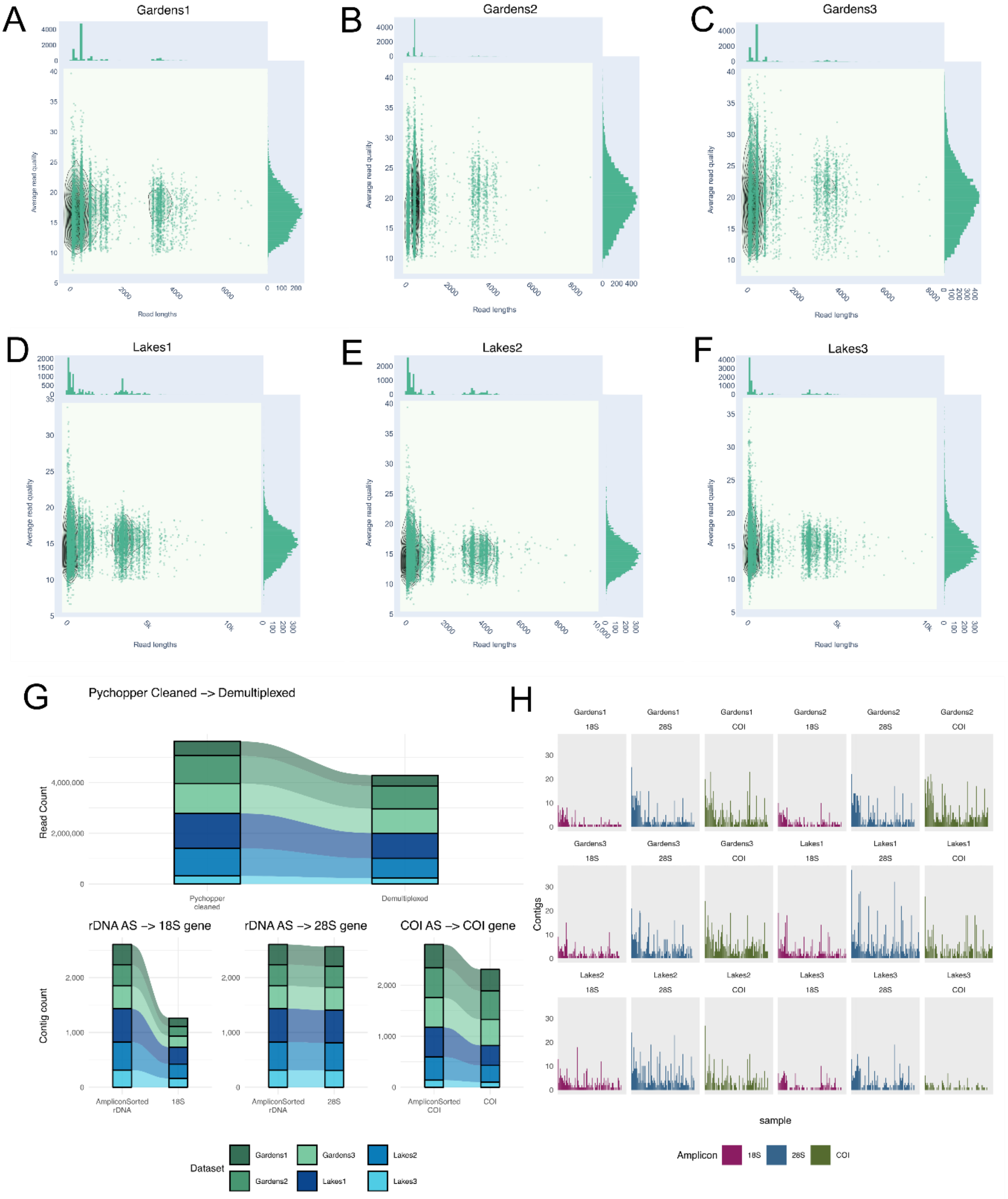

**Supplementary Fig. 2:** Quality and counts of reads and contigs across pipeline stages. (A-F) Kernel density plots with histograms of read quality scores along the Y-axis and read lengths along the X-axis. Each point is one read, and histograms show overall distribution of metrics Plots show datasets Gardens 1, Gardens 2, Gardens 3, Lakes 1, Lakes 2, and Lakes 3, respectively (G) Panelled alluvial plots showing flow of read counts and contig counts across bioinformatic processing stages (H) Panelled bar plots of counts of contigs (Y-axis) per dataset per gene amplified (X-axis).

#### Supplementary Figure 3

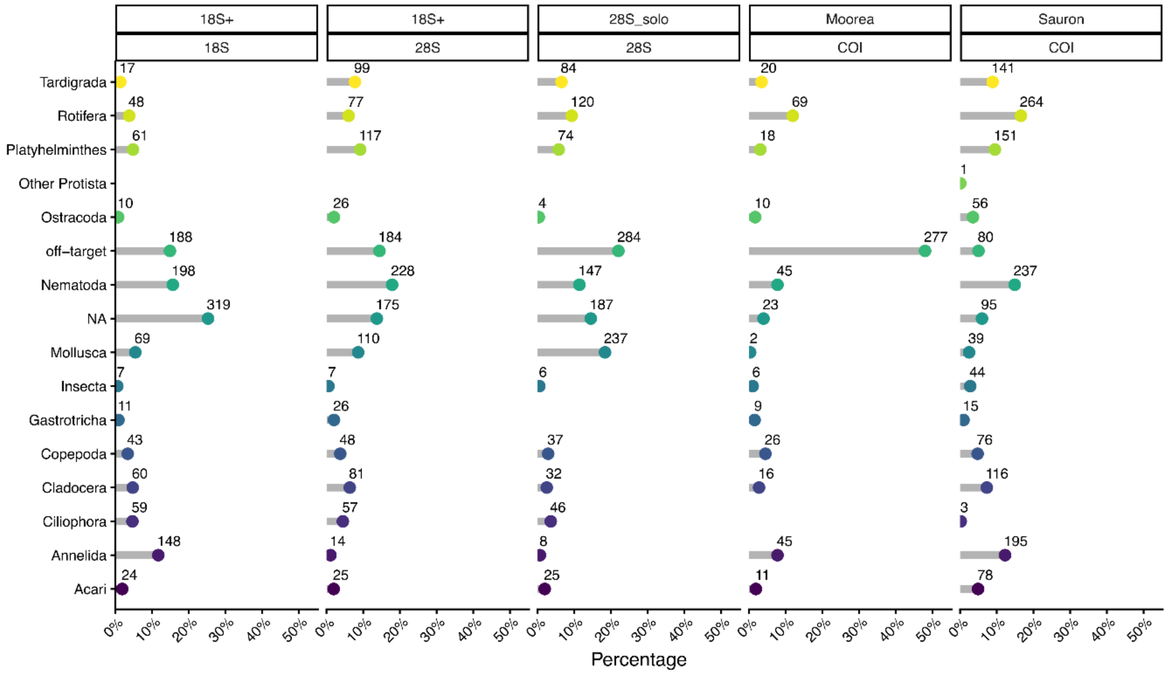

**Supplementary Fig. 3:** Taxonomic assignment with automated blastn and LCA approach for all contigs. Lollipop plot with intentionally sampled higher taxa on the y axis and counts of contigs per taxon on the x-axis, panelled by amplicon

### Supplementary Figure 4

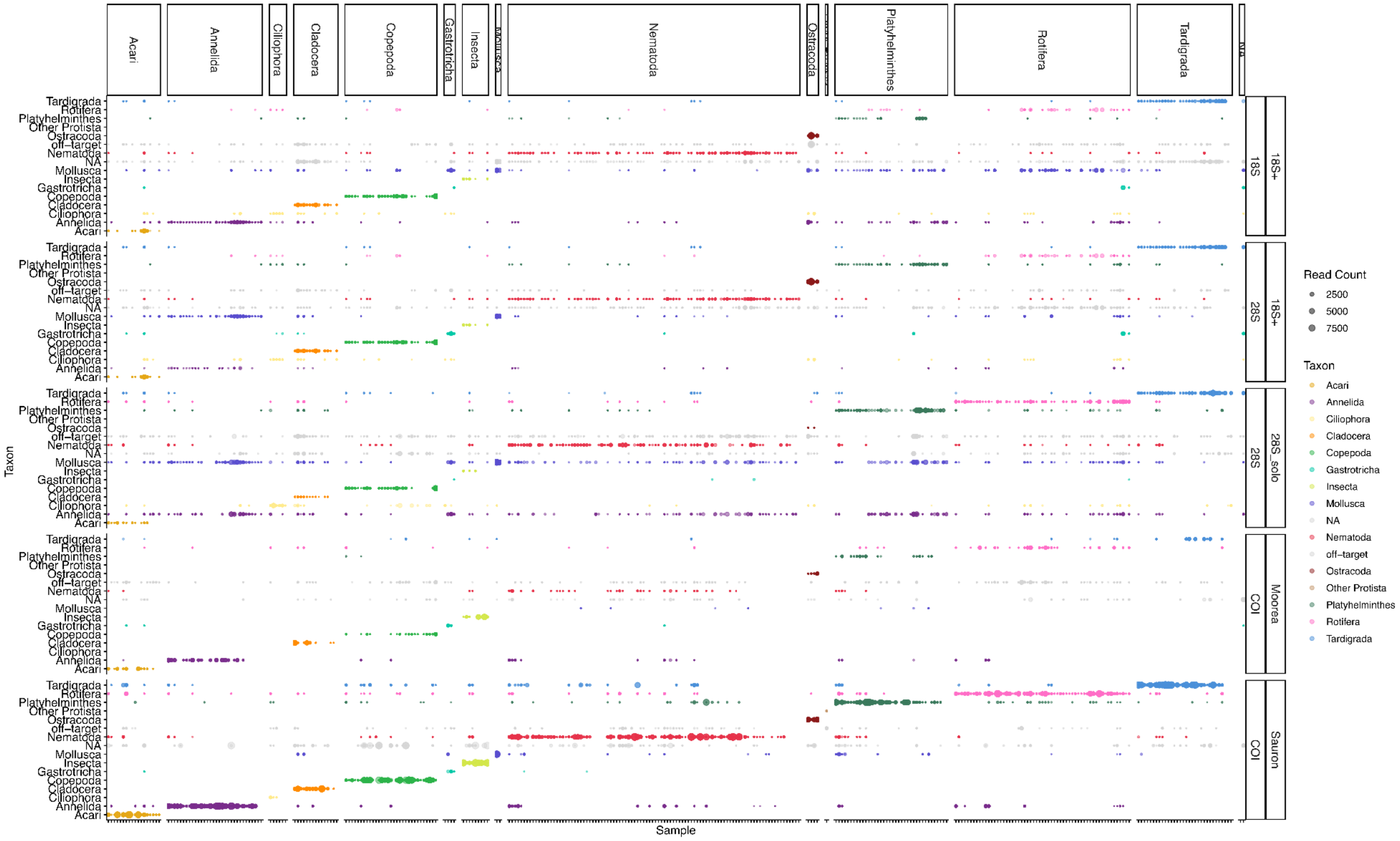

**Supplementary Fig. 4**: Bubble plot of showing LCA of each contig panelled per sample per barcode, at broader taxonomic level. Samples unlabelled on the X axis, grouped by intended taxon sampled. Bubbles sized by read counts per contig.

### Supplementary Tables

#### Supplementary Table 1

| *Locus* | Barcode ID | Forward Primer ID | Forward Sequence (5'-3') | Forward Tm (°C) | Forward Source | Reverse Primer ID | Reverse Primer Sequence (5'-3') | Reverse Tm (°C) | Reverse Primer Source | PCR conditions | Amplicon Size (bp) |
|---|---|---|---|---|---|---|---|---|---|---|---|
| *rDNA* | 18S+ | SSU_F04 | [TGTAAAACGACGGCCAG] GCTTGTCTCAAAGATTAAGC*C | 65.9 | Blaxter et al., 1998 | 28S_3RC | [CAGGAAACAGCTATGAC] CRCCAGTTCTGCTTACCAAA*A | 65.2 | Machida and Knowlton, 2012 | 1x[98°C/2:00], 35x[98°C/0:10, 62°C/0:15, 68°C/6:00], 1x[68°C/5:00] | 4000-5000 |
| *rDNA* | 28S | F63.2 | [TGTAAAACGACGGCCAG] ACCCGCTGAAYTTAAGCATA*T | 66 | Passamaneck et al., 2004 | R3264.2 | [CAGGAAACAGCTATGAC] TWCYRMCTTAGAGGCGTTCA*G | 64.5 | Passamaneck et al., 2004 | 1x[98°C/2:00], 35x[98°C/0:10, 62°C/0:15, 68°C/6:00], 1x[68°C/5:00] | 3200 |
| *COI* | Sauron | Sauron-S878 | [TGTAAAACGACGGCCAG] GGDRCWGGWTGAACWGTWTAYCCNC*C | 69.3 | Rennstam Rubbmark et al., 2018 | jgHCO2198 | [CAGGAAACAGCTATGAC] TAIACYTCIGGRTGICCRAARAAYC*A | 67.9 | Geller et al., 2013 | 1x[94°C/2:00], 40x[94°C/0:15, 48°C/0:15, 68°C/1:00], 1x[68°C/5:00] | 313 |
| *COI* | Moorea | jgLCO1490 | [TGTAAAACGACGGCCAG] TITCIACIAAYCAYAARGAYATTG*G | 66.9 | Geller et al., 2013 | jgHCO2198 | [CAGGAAACAGCTATGAC] TAIACYTCIGGRTGICCRAARAAYC*A | 67.9 | Geller et al., 2013 | 1x[94°C/2:00], 40x[94°C/0:15, 48°C/0:15, 68°C/1:00], 1x[68°C/5:00] | 658 |
| *Indexing* | Indexing | SP5_fb17.2_001_M13F | {CATGTAATGCACGTACTTTCAGGGT} GAGCGTCTAATCGTAAT [TGTAAAACGACGGCCA*G] | | | SP27_fb17.2_001_M13R | {GATCAGGTGAGGCTGCGACGACT} CCTCCGTGCCTGGTTAA [CAGGAAACAGCTATGA*C] | | | | |
| *Indexing* | Indexing | SP5_fb17.2_002_M13F | {CATGTAATGCACGTACTTTCAGGGT} CTACCGTGGATATTCAA [TGTAAAACGACGGCCA*G] | | | SP27_fb17.2_002_M13R | {GATCAGGTGAGGCTGCGACGACT} AACTTCAGGTCCACAGC [CAGGAAACAGCTATGA*C] | | | | |
| *Indexing* | Indexing | SP5_fb17.2_003_M13F | {CATGTAATGCACGTACTTTCAGGGT} AATTCCACTTACAACGG [TGTAAAACGACGGCCA*G] | | | SP27_fb17.2_003_M13R | {GATCAGGTGAGGCTGCGACGACT} ACGCGGTGGTGTAACGA [CAGGAAACAGCTATGA*C] | | | | |
| *Indexing* | Indexing | SP5_fb17.2_004_M13F | {CATGTAATGCACGTACTTTCAGGGT} AGTGTGCCGCCAACCAA [TGTAAAACGACGGCCA*G] | | | SP27_fb17.2_004_M13R | {GATCAGGTGAGGCTGCGACGACT} AAGAATGGATAAGGAGG [CAGGAAACAGCTATGA*C] | | | | |
| *Indexing* | Indexing | SP5_fb17.2_005_M13F | {CATGTAATGCACGTACTTTCAGGGT} AGCCTCATTGGTTGTTC [TGTAAAACGACGGCCA*G] | | | SP27_fb17.2_005_M13R | {GATCAGGTGAGGCTGCGACGACT} ATAGGTCATTGCGCTTC [CAGGAAACAGCTATGA*C] | | | | |
| *Indexing* | Indexing | SP5_fb17.2_006_M13F | {CATGTAATGCACGTACTTTCAGGGT} GATTCTACAAGTGGTGA [TGTAAAACGACGGCCA*G] | | | SP27_fb17.2_006_M13R | {GATCAGGTGAGGCTGCGACGACT} CCGATCCTTCAGAGCCA [CAGGAAACAGCTATGA*C] | | | | |
| *Indexing* | Indexing | SP5_fb17.2_007_M13F | {CATGTAATGCACGTACTTTCAGGGT} ACAGGTTGCCGGAGTCT [TGTAAAACGACGGCCA*G] | | | SP27_fb17.2_007_M13R | {GATCAGGTGAGGCTGCGACGACT} CGCTGCTAGAATATGCC [CAGGAAACAGCTATGA*C] | | | | |
| *Indexing* | Indexing | SP5_fb17.2_008_M13F | {CATGTAATGCACGTACTTTCAGGGT} CAATCGTGACCATCCGG [TGTAAAACGACGGCCA*G] | | | SP27_fb17.2_008_M13R | {GATCAGGTGAGGCTGCGACGACT} ATTGGACTGTTAGGAGG [CAGGAAACAGCTATGA*C] | | | | |
| *Indexing* | Indexing | SP5_fb17.2_009_M13F | {CATGTAATGCACGTACTTTCAGGGT} AACAACAACAACAACCG [TGTAAAACGACGGCCA*G] | | | | | | | | |
| *Indexing* | Indexing | SP5_fb17.2_010_M13F | {CATGTAATGCACGTACTTTCAGGGT} GGTCAGGTAGTCCGTAT [TGTAAAACGACGGCCA*G] | | | | | | | | |
| *Indexing* | Indexing | SP5_fb17.2_011_M13F | {CATGTAATGCACGTACTTTCAGGGT} GCCTGTGCGGAGTAGAT [TGTAAAACGACGGCCA*G] | | | | | | | | |
| *Indexing* | Indexing | SP5_fb17.2_012_M13F | {CATGTAATGCACGTACTTTCAGGGT} CCAACGGACTACGAATT [TGTAAAACGACGGCCA*G] | | | | | | | | |

**Supplementary Table 1:** Primer information for OrCA-seq. Square brackets indicate inclusion of the M13 sequence. Tm's taken from manufacturers recommendation. Indexing primers follow this format: {fixed region} index [M13 tag]. Indexing PCR conditions: 1x[94°C/2:00], 7x[94°C/0:15, 47°C/0:15, 68°C/4:00], 1x[68°C/5:00]. Indexing primers are all 5' Phosphorylated. The indexing primers (12 forward and 8 reverse) are used in a dual-combination strategy to enable each of the 96 samples in the plate to have a unique pair of index primers.

### Supplementary Table 2

| ***Dataset*** | **Date of run** | **Flow Cell Type** | **Flow Cell ID** | **Run Length (hh:mm)** | **Passed Output (Gb)** | **Passed Reads (M)** | **N50 (kb)** |
|---|---|---|---|---|---|---|---|
| *Gardens1* | 15/08/2025 | FLO-MIN114 | FAY74148 | 8:00 | 1.32 | 1.22 | 3.3 |
| *Gardens2* | 16/08/2025 | FLO-PRO114M | PAY14142 | 1:47 | 1.90 | 1.87 | 2.12 |
| *Gardens3* | 16/08/2025 | FLO-PRO114M | PAW88913 | 1:46 | 2.51 | 2.07 | 3.48 |
| *Lakes1* | 07/09/2025 | FLO-MIN114 | FAY75466 | 13:39 | 4.62 | 2.61 | 3.58 |
| *Lakes2* | 08/09/2025 | FLO-MIN114 | FAY75466 | 14:00 | 3.07 | 2.10 | 3.63 |
| *Lakes3* | 09/09/2025 | FLO-MIN114 | FAY75466 | 15:37 | 0.89 | 0.74 | 3.56 |

**Supplementary Table 2:** MinKNOW run statistics from ORCA-seq sequencing runs.

#### Supplementary Table 3

| *Gene* | Taxa | Species Name | Genbank Acc. # |
|---|---|---|---|
| *COI* | Platyhelminthes | Astrotorhynchus bifidus | MG256167.1 |
| *COI* | Annelida | Mesenchytraeus beumeri | PP181880.1 |
| *COI* | Nematoda | Necator americanus | MH200973.1 |
| *COI* | Rotifera | Adineta vaga | DQ079961.1 |
| *COI* | Tardigrada | Milnesium tardigradum | OP009210.1 |
| *COI* | Gastrotricha | Chaetonotus acanthodes | MN493713.1 |
| *COI* | Acari | Tydeus sp. | HQ927816.1 |
| *COI* | Ciliophora | Paramecium bursaria | OK356534.1 |
| *COI* | Copepoda | Phyllognathopus viguieri | KP845501.1 |
| *COI* | Cladocera | Daphnia Magna | GU680598.1 |
| *18S* | Platyhelminthes | Astrotorhynchus bifidus | MG256065.1 |
| *18S* | Annelida | Chamaedrilus sphagnetorum | KX618777.1 |
| *18S* | Nematoda | C. elegans | NR_000054.1 |
| *18S* | Rotifera | Adineta vaga | KM043254.2 |
| *18S* | Tardigrada | Milnesium tardigradum | HM187581.1 |
| *18S* | Gastrotricha | Chaetonotus acanthodes | JQ798585.1 |
| *18S* | Acari | Thalassozetes grenadensis | MZ220310.1 |
| *18S* | Ciliophora | Paramecium bursaria | MG589318.1 |
| *18S* | Copepoda | Volkmannia attenuata | MF077776.1 |
| *18S* | Cladocera | Scapholeberis rammneri | KM244570.1 |
| *28S* | Platyhelminthes | Astrotorhynchus bifidus | MG256110.1 |
| *28S* | Annelida | Diplocardia conoyeri | HQ728983.1 |
| *28S* | Nematoda | Xiphinema rivesi | AY210845.1 |
| *28S* | Rotifera | Adineta vaga | DQ089739.1 |
| *28S* | Tardigrada | Milnesium tardigradum | FJ435780.1 |
| *28S* | Gastrotricha | Chaetonotus acanthodes | JQ798653.1 |
| *28S* | Acari | Odontocepheus oblongus | KY922089.1 |
| *28S* | Ciliophora | Euplotes aediculatus | AF223571.1 |
| *28S* | Copepoda | Salmincola edwardsi | DQ180346.2 |
| *28S* | Cladocera | Daphnia magna | EU370436.1 |

**Supplementary Table 3:** Genbank sequences used to screen contig files for the presence of expected taxa in alternate contigs, as a part of the manual BLAST curation.

#### Supplementary Table 4

| *Taxon & Gene* | Alignment algorithm | Whitelist divergence threshold (%) | Deduplication similarity threshold (%) | Overlap threshold between samples and references (bp) |
|---|---|---|---|---|
| *Platyhelminthes 28S* | E-INS-I | 96 | 2 | 0 |
| *Annelida COI* | L-INS-I | 96 | 0 (no dedup) | 300 |
| *Nematoda 18S* | E-INS-I | 96 | 2 | 0 |

**Supplementary Table 4:** Phylogenetic tree reconstruction metrics.

### Supplementary Table 5

| ***Barcode and Primer*** | **Min Readcount** | **Q1 Readcount** | **Median Readcount** | **Mean Readcount** | **Q3 Readcount** | **Max Readcount** |
|---|---|---|---|---|---|---|
| *18S , 18S+* | 5 | 23 | 53 | 229.58 | 171.75 | 8706 |
| *28S , 18S+* | 5 | 23 | 54 | 228.39 | 171 | 8706 |
| *28S , 28S_solo* | 5 | 29 | 67 | 291.68 | 220 | 7332 |
| *COI , Moorea* | 5 | 17 | 44 | 214.14 | 129 | 5859 |
| *COI , Sauron* | 4 | 40 | 113 | 649.14 | 428.5 | 9862 |

**Supplementary Table 5:** Read count statistics for assembled amplicon contigs.

### Supplementary Table 6

| Manual BLAST outcome | Taxa | n | Taxa-relative percentage | Cumulative outcome percentage |
|---|---|---|---|---|
| No contig present | **Acari** | 32 | 23.88 | |
| Off target/No Match | **Acari** | 33 | 24.63 | 48.51 |
| AC BLAST match | **Acari** | 4 | 2.99 | |
| MRC BLAST match | **Acari** | 65 | 48.51 | 51.49 |
| No contig present | **Annelida** | 39 | 22.94 | |
| Off target/No Match | **Annelida** | 15 | 8.82 | 31.76 |
| AC BLAST match | **Annelida** | 6 | 3.53 | |
| MRC BLAST match | **Annelida** | 110 | 64.71 | 68.24 |
| No contig present | **Ciliophora** | 17 | 53.13 | |
| Off target/No Match | **Ciliophora** | 4 | 12.50 | 65.63 |
| AC BLAST match | **Ciliophora** | 3 | 9.38 | |
| MRC BLAST match | **Ciliophora** | 8 | 25.00 | 34.38 |
| No contig present | **Cladocera** | 62 | 31.63 | |
| Off target/No Match | **Cladocera** | 18 | 9.18 | 40.82 |
| AC BLAST match | **Cladocera** | 23 | 11.73 | |
| MRC BLAST match | **Cladocera** | 93 | 47.45 | 59.18 |
| No contig present | **Copepoda** | 41 | 23.84 | |
| Off target/No Match | **Copepoda** | 28 | 16.28 | 40.12 |
| AC BLAST match | **Copepoda** | 12 | 6.98 | |
| MRC BLAST match | **Copepoda** | 91 | 52.91 | 59.88 |
| No contig present | **Gastrotricha** | 7 | 14.58 | |
| Off target/No Match | **Gastrotricha** | 15 | 31.25 | 45.83 |
| AC BLAST match | **Gastrotricha** | 1 | 2.08 | |
| MRC BLAST match | **Gastrotricha** | 25 | 52.08 | 54.17 |
| No contig present | **Nematoda** | 200 | 34.01 | |
| Off target/No Match | **Nematoda** | 129 | 21.94 | 55.95 |
| AC BLAST match | **Nematoda** | 20 | 3.40 | |
| MRC BLAST match | **Nematoda** | 239 | 40.65 | 44.05 |
| No contig present | **Ostracoda** | 13 | 19.12 | |
| Off target/No Match | **Ostracoda** | 5 | 7.35 | 26.47 |
| AC BLAST match | **Ostracoda** | 5 | 7.35 | |
| MRC BLAST match | **Ostracoda** | 45 | 66.18 | 73.53 |
| No contig present | **Platyhelminthes** | 56 | 25.00 | |
| Off target/No Match | **Platyhelminthes** | 63 | 28.13 | 53.13 |
| AC BLAST match | **Platyhelminthes** | 9 | 4.02 | |
| MRC BLAST match | **Platyhelminthes** | 96 | 42.86 | 46.88 |
| No contig present | **Rotifera** | 59 | 18.44 | |
| Off target/No Match | **Rotifera** | 81 | 25.31 | 43.75 |
| AC BLAST match | **Rotifera** | 21 | 6.56 | |
| MRC BLAST match | **Rotifera** | 159 | 49.69 | 56.25 |
| No contig present | **Tardigrada** | 56 | 28.00 | |
| Off target/No Match | **Tardigrada** | 33 | 16.50 | 44.50 |
| AC BLAST match | **Tardigrada** | 11 | 5.50 | |
| MRC BLAST match | **Tardigrada** | 100 | 50.00 | 55.50 |

**Supplementary Table 6:** Outcomes of manual BLAST curation per taxa group. The fourth column contains relative percentages of all four possible outcomes within the taxonomic group. whilst the fifth column contains cumulative percentages relating to barcode recovery. Row shading indicates recovery (white), or non-recovery (grey) of barcodes. Recovered barcodes originate from either an Alternate Contig (AC), or a Max Readcount Contig (MRC).

#### Supplementary Table 7.A

| *Gene* | Expected Taxon | Contig Origin/ Result type | n | Gene-Taxon percent. | Recovered contig percent. |
|---|---|---|---|---|---|
| *18S* | Acari | No contig present | 11 | 33.33 | |
| *18S* | Acari | Off target/No Match | 6 | 18.18 | |
| *18S* | Acari | 18S+ | 16 | 48.48 | 100.00 |
| *18S* | Annelida | No contig present | 6 | 13.95 | |
| *18S* | Annelida | Off target/No Match | 4 | 9.30 | |
| *18S* | Annelida | 18S+ | 33 | 76.74 | 100.00 |
| *18S* | Ciliophora | No contig present | 3 | 37.50 | |
| *18S* | Ciliophora | 18S+ | 5 | 62.50 | 100.00 |
| *18S* | Cladocera | No contig present | 10 | 20.41 | |
| *18S* | Cladocera | Off target/No Match | 3 | 6.12 | |
| *18S* | Cladocera | 18S+ | 36 | 73.47 | 100.00 |
| *18S* | Copepoda | No contig present | 9 | 20.93 | |
| *18S* | Copepoda | Off target/No Match | 2 | 4.65 | |
| *18S* | Copepoda | 18S+ | 32 | 74.42 | 100.00 |
| *18S* | Gastrotricha | Off target/No Match | 5 | 41.67 | |
| *18S* | Gastrotricha | 18S+ | 7 | 58.33 | 100.00 |
| *18S* | Nematoda | No contig present | 37 | 25.17 | |
| *18S* | Nematoda | Off target/No Match | 18 | 12.24 | |
| *18S* | Nematoda | 18S+ | 92 | 62.59 | 100.00 |
| *18S* | Ostracoda | No contig present | 1 | 5.88 | |
| *18S* | Ostracoda | 18S+ | 16 | 94.12 | 100.00 |
| *18S* | Platyhelminthes | No contig present | 10 | 17.86 | |
| *18S* | Platyhelminthes | Off target/No Match | 3 | 5.36 | |
| *18S* | Platyhelminthes | 18S+ | 43 | 76.79 | 100.00 |
| *18S* | Rotifera | No contig present | 22 | 27.50 | |
| *18S* | Rotifera | Off target/No Match | 30 | 37.50 | |
| *18S* | Rotifera | 18S+ | 28 | 35.00 | 100.00 |
| *18S* | Tardigrada | No contig present | 9 | 18.00 | |
| *18S* | Tardigrada | Off target/No Match | 6 | 12.00 | |
| *18S* | Tardigrada | 18S+ | 35 | 70.00 | 100.00 |

#### Supplementary Table 7.B

| *Gene* | Expected Taxon | Contig Origin/ Result type | n | Gene-Taxon percent. | Recovered contig percent. |
|---|---|---|---|---|---|
| *28S* | Acari | No contig present | 6 | 18.18 | |
| *28S* | Acari | Off target/No Match | 6 | 18.18 | |
| *28S* | Acari | 18S+ | 11 | 33.33 | 52.38 |
| *28S* | Acari | 28S | 10 | 30.30 | 47.62 |
| *28S* | Annelida | No contig present | 6 | 13.95 | |
| *28S* | Annelida | Off target/No Match | 4 | 9.30 | |
| *28S* | Annelida | 18S+ | 7 | 16.28 | 21.21 |
| *28S* | Annelida | 28S | 26 | 60.47 | 78.79 |
| *28S* | Ciliophora | No contig present | 2 | 25.00 | |
| *28S* | Ciliophora | 18S+ | 1 | 12.50 | 16.67 |
| *28S* | Ciliophora | 28S | 5 | 62.50 | 83.33 |
| *28S* | Cladocera | No contig present | 8 | 16.33 | |
| *28S* | Cladocera | Off target/No Match | 5 | 10.20 | |
| *28S* | Cladocera | 18S+ | 35 | 71.43 | 97.22 |
| *28S* | Cladocera | 28S | 1 | 2.04 | 2.78 |
| *28S* | Copepoda | No contig present | 7 | 16.28 | |
| *28S* | Copepoda | Off target/No Match | 6 | 13.95 | |
| *28S* | Copepoda | 18S+ | 19 | 44.19 | 63.33 |
| *28S* | Copepoda | 28S | 11 | 25.58 | 36.67 |
| *28S* | Gastrotricha | Off target/No Match | 4 | 33.33 | |
| *28S* | Gastrotricha | 18S+ | 4 | 33.33 | 50.00 |
| *28S* | Gastrotricha | 28S | 4 | 33.33 | 50.00 |
| *28S* | Nematoda | No contig present | 23 | 15.65 | |
| *28S* | Nematoda | Off target/No Match | 9 | 6.12 | |
| *28S* | Nematoda | 18S+ | 31 | 21.09 | 26.96 |
| *28S* | Nematoda | 28S | 84 | 57.14 | 73.04 |
| *28S* | Ostracoda | Off target/No Match | 2 | 11.76 | |
| *28S* | Ostracoda | 18S+ | 13 | 76.47 | 86.67 |
| *28S* | Ostracoda | 28S | 2 | 11.76 | 13.33 |
| *28S* | Platyhelminthes | No contig present | 5 | 8.93 | |
| *28S* | Platyhelminthes | Off target/No Match | 1 | 1.79 | |
| *28S* | Platyhelminthes | 18S+ | 1 | 1.79 | 2.00 |
| *28S* | Platyhelminthes | 28S | 49 | 87.50 | 98.00 |
| *28S* | Rotifera | No contig present | 7 | 8.75 | |
| *28S* | Rotifera | Off target/No Match | 10 | 12.50 | |
| *28S* | Rotifera | 18S+ | 14 | 17.50 | 22.22 |
| *28S* | Rotifera | 28S | 49 | 61.25 | 77.78 |
| *28S* | Tardigrada | No contig present | 4 | 8.00 | |
| *28S* | Tardigrada | Off target/No Match | 7 | 14.00 | |
| *28S* | Tardigrada | 18S+ | 9 | 18.00 | 23.08 |
| *28S* | Tardigrada | 28S | 30 | 60.00 | 76.92 |

Supplementary Table 7.C

| *Gene* | Expected Taxon | Contig Origin/ Result type | n | Gene-Taxon percent. | Recovered contig percent. |
|---|---|---|---|---|---|
| *COI* | Acari | No contig present | 15 | 22.39 | |
| *COI* | Acari | Off target/No Match | 21 | 31.34 | |
| *COI* | Acari | Moorea | 8 | 11.94 | 25.81 |
| *COI* | Acari | Sauron | 23 | 34.33 | 74.19 |
| *COI* | Annelida | No contig present | 27 | 32.14 | |
| *COI* | Annelida | Off target/No Match | 7 | 8.33 | |
| *COI* | Annelida | Moorea | 18 | 21.43 | 36.00 |
| *COI* | Annelida | Sauron | 32 | 38.10 | 64.00 |
| *COI* | Ciliophora | No contig present | 12 | 75.00 | |
| *COI* | Ciliophora | Off target/No Match | 4 | 25.00 | |
| *COI* | Cladocera | No contig present | 44 | 44.90 | |
| *COI* | Cladocera | Off target/No Match | 10 | 10.20 | |
| *COI* | Cladocera | Moorea | 9 | 9.18 | 20.45 |
| *COI* | Cladocera | Sauron | 35 | 35.71 | 79.55 |
| *COI* | Copepoda | No contig present | 25 | 29.07 | |
| *COI* | Copepoda | Off target/No Match | 20 | 23.26 | |
| *COI* | Copepoda | Moorea | 13 | 15.12 | 31.71 |
| *COI* | Copepoda | Sauron | 28 | 32.56 | 68.29 |
| *COI* | Gastrotricha | No contig present | 7 | 29.17 | |
| *COI* | Gastrotricha | Off target/No Match | 6 | 25.00 | |
| *COI* | Gastrotricha | Moorea | 5 | 20.83 | 45.45 |
| *COI* | Gastrotricha | Sauron | 6 | 25.00 | 54.55 |
| *COI* | Nematoda | No contig present | 140 | 47.62 | |
| *COI* | Nematoda | Off target/No Match | 102 | 34.69 | |
| *COI* | Nematoda | Moorea | 16 | 5.44 | 30.77 |
| *COI* | Nematoda | Sauron | 36 | 12.24 | 69.23 |
| *COI* | Ostracoda | No contig present | 12 | 35.29 | |
| *COI* | Ostracoda | Off target/No Match | 3 | 8.82 | |
| *COI* | Ostracoda | Moorea | 6 | 17.65 | 31.58 |
| *COI* | Ostracoda | Sauron | 13 | 38.24 | 68.42 |
| *COI* | Platyhelminthes | No contig present | 41 | 36.61 | |
| *COI* | Platyhelminthes | Off target/No Match | 59 | 52.68 | |
| *COI* | Platyhelminthes | Moorea | 3 | 2.68 | 25.00 |
| *COI* | Platyhelminthes | Sauron | 9 | 8.04 | 75.00 |
| *COI* | Rotifera | No contig present | 30 | 18.75 | |
| *COI* | Rotifera | Off target/No Match | 41 | 25.63 | |
| *COI* | Rotifera | Moorea | 25 | 15.63 | 28.09 |
| *COI* | Rotifera | Sauron | 64 | 40.00 | 71.91 |
| *COI* | Tardigrada | No contig present | 43 | 43.00 | |
| *COI* | Tardigrada | Off target/No Match | 20 | 20.00 | |
| *COI* | Tardigrada | Moorea | 8 | 8.00 | 21.62 |
| *COI* | Tardigrada | Sauron | 29 | 29.00 | 78.38 |

**Supplemental Table 7:** Recovery percentages for each taxon. Split into 18S (7.A), 28S (7.B), COI (7.C). Row shading indicates recovery (white), or non-recovery (grey) of barcodes. Within the Contig origin/result type, any white shaded rows record the amplicon of origin. Gene-Taxon percent. is the percentage of each result type within the taxa group. Multiple different amplicons were generated for both 28S and COI. The proportion between the successfully recovered amplicons is shown as a percentage in the final column.

### Supplemental Table 8

| Assignment | Taxon | Gene | Type | Top BLASTn %ID hit | Top BLASTn %ID | LCA | Note |
|---|---|---|---|---|---|---|---|
| Nais SP275SP56 L2 | Annelida | *COI* | Sample, Lakes | Pristina longiseta complex PP136959.1 | 91.96 | Pristina genus | |
| Ripistes SP277SP56 L2 | Annelida | *COI* | Sample, Lakes | Pristina longiseta complex PP136959.1 | 91.96 | Pristina genus | |
| Vejdovskyella comata SP276SP56 L2 | Annelida | *COI* | Sample, Lakes | Ripistes parasita KY633403.1 | 100.00 | Naidinae subfamily | |
| Vejdovskyella comata SP274SP56 L2 | Annelida | *COI* | Sample, Lakes | Specaria josinae voucher PP137100.1 | 100.00 | Specaria josinae | |
| Marionina SP272SP55 L1 | Annelida | *COI* | Sample, Lakes | Enchytronia parva complex PP180780.1 | 98.78 | Enchytronia parva | |
| Henlea SP273SP55 L1 | Annelida | *COI* | Sample, Lakes | Fridericia perrieri complex PP181205.1 | 99.68 | Fridericia perrieri | |
| HM386184.1 Haplotaxida sp. | Annelida | *COI* | Reference | Fridericia connata PP180973.1 | 86.91 | Fridericia connata (exc. Other Haplotaxida) | Phylogenetic placement incorrect. Second best BLAST hit noted. |
| Enchytraeus SP271SP55 L1 | Annelida | *COI* | Sample, Lakes | Achaeta unibulba PP180120.1 | 100.00 | Achaeta unibulba | |
| Punctodora ratzeburgensis SP276SP511 L2 | Nematoda | *18S* | Sample, Lakes | Eumonhystera cf. hungarica KJ636237.1 | 98.64 | Monhysteridae family (exc. Uncultured eukaryotes) | |
| Theristus agilis SP27 005 SP5 011 day3 | Nematoda | *18S* | Sample, Lakes | Geomonhystera sp KJ636213.1 | 96.07 | Monhysteridae family | |
| Theristus agilis SP27 006 SP5 011 day3 | Nematoda | *18S* | Sample, Lakes | Geomonhystera sp KJ636213.1 | 96.07 | Monhysteridae family | |
| Mermithidae SP271SP511 L3 | Nematoda | *18S* | Sample, Lakes | Aporcelaimellus sp AJ875154.1 | 99.94 | Aporcelaimellus (exc. Uncultured apicomplexan) | |
| Prionchulus muscorum SP275SP57 L3 | Nematoda | *18S* | Sample, Lakes | Mylonchulus brachyuris AB361437.1 | 99.02 | Mylonchulus genus | |
| Prionchulus muscorum SP276SP57 L3 | Nematoda | *18S* | Sample, Lakes | Mylonchulus brachyuris AB361437.1 | 99.02 | Mylonchulus genus | |
| Achromadora ruricola SP277SP56 L3 | Nematoda | *18S* | Sample, Lakes | Ethmolaimus pratensis AY593942.1 | 99.82 | Cyatholaimidae family | |
| Epidorilaimus centrocercus SP272SP57 L3 | Nematoda | *18S* | Sample, Lakes | Mesodorylaimus cf. centrocercus | - | - | Mislabelled at sampling stage |

Supplemental Table 8: Misidentified tips in phylogenetic trees. Columns note tip assignment, taxon, gene, tip dataset or reference, BLASTn 'best hit' according to percentage identity and accession, percent identity for that BLASTn hit, Lowest common ancestor of top 5 hits based on percent identity, and relevant notes.
